# Dynamics of living cells in a cytomorphological state space

**DOI:** 10.1101/549246

**Authors:** Amy Y. Chang, Wallace F. Marshall

## Abstract

Cells are non-equilibrium systems that rely on a continuous exchange of matter and energy with the environment to sustain their metabolic needs. The non-equilibrium nature of this system presents considerable challenges to developing a general theory describing its behavior; however, studies have demonstrated that when studied at appropriate spatiotemporal scales, the behavior of ensembles of non-equilibrium systems can resemble that of system at equilibrium. Here we apply this principle to a population of cells within a cytomorphological state space and demonstrate that cellular transition dynamics within this space can be suitably described using equilibrium dynamics formalisms. We use this framework to map the effective energy landscape underlying the cytomorphological state space of a population of mouse embryonic fibroblasts (MEFs) and identify topographical non-uniformity in this space, indicating non-uniform occupation of cytomorphological states within an isogenic population. The introduction of exogenous apoptotic agents altered this energy landscape, inducing formation of additional energy minima that correlated directly with changes in sensitivity to apoptotic induction. The measured application of equilibrium dynamics formalism allows us to accurately capture and these findings suggest that though cells are complex non-equilibrium systems, the application of formalisms derived from equilibrium thermodynamics can provide insight into the basis of non-genetic heterogeneities as well as the relationship between morphological and functional heterogeneity.

## Main Text

Cells are non-equilibrium systems that continuously exchange matter and energy with their environments, encapsulate a range of irreversible chemical reactions, and exhibit time-dependent fluctuations in the concentration of their DNA, mRNA, proteins, and other biomolecules. Given the central importance of cells in biological systems, significant efforts over the years have been directed at developing theorems to describe this system; however, a unifying theory remains, at present, out of reach. In recent years, several studies have indicated that when viewed at appropriate spatiotemporal scales, the behavior of an ensemble of non-equilibrium systems can at times be accurately approximated using equilibrium dynamics formalisms (*1-4*). Here, we demonstrate the applicability of this principle in a population of isogenic mouse embryonic fibroblasts (MEFs) and apply an equilibrium dynamics-based framework to uncover a new basis for functional heterogeneity within isogenic populations.

We begin by identifying an appropriate parameter space that would allow both for testing the applicability of an equilibrium dynamics framework and for tracking the relationship between state space dynamics and functional behavior in a population of living cells. The state space of a living cell can be defined using many different parameter sets. The set relating most directly to cellular function is the molecular microstate, which represents the sum of the epigenetic, transcriptomic, proteomic, and additional molecular states of the cell. The ensemble of these states define the functional state of a cell; consequently, variability in the molecular microstate can lead directly to functional heterogeneity within isogenic populations. This phenomenon is known as non-genetic heterogeneity, wherein isogenic cells exhibit extensive functional variability; these non-genetic heterogeneities, play important roles across a diverse range of biological processes (*5,6*). In mammalian cells, these include the selective differentiation of hematopoietic progenitor cells (*7*), while in bacterial cells, these include the appearance of subpopulations of “persister” cells that are preferentially able to survive antibiotic exposure (*8*). Non-genetic heterogeneities remain of central interest in cell biology, with many of the underlying laws governing this phenomenon remaining unknown.

In order to explore the applicability of equilibrium dynamics formalisms in studying non-genetic heterogeneity, we first identified methods for approximating the molecular state of living cells over an experimentally relevant time course. Whole-genome methods (e.g. transcriptomics, proteomics, ChiP-Seq) offer by far the most comprehensive catalogue of the microstate (*9-14*), but these methods are cell-destructive in nature and of limited use in live cell experiments. Fluorescent reporters linked to mRNA and protein targets can be tracked over multi-hour time courses in living cells (*15,16*); however, reporters are limited to a few target sequences at a time, offering a limited snapshot of the molecular microstate. Here we employ cytomorphology, as defined at the cell and organelle scale, as a proxy for the molecular microstate. Cytomorphology represents the sum of many thousands of molecular processes (*17-19*), providing global cell state information that offers a compromise between experimental accessibility and a comprehensive catalog of the molecular microstate. Further underlining the link between the cytomorphological and functional states of the cell are numerous studies demonstrating that diseases such as cancers (*20*) and neurodegenerative disorders (*21,22*) are characterized by concomitant functional and cytomorphological changes. By altering the scale at which we define cellular state, we developed methods that allowed us to track state space occupancy in living cells and identify relationships between state space dynamics of the population as a whole and functional heterogeneity.

We began by developing a methodology to measure cytomorphological features and quantify state space occupancy in a population of cells. We chose mouse embryonic fibroblasts (MEFs) as our model population, as Wild-type MEFs (WT MEFs) exhibit native cytomorphological heterogeneity, rendering them ideal candidates for studying non-genetic heterogeneity in the context of cytomorphological variability. WT MEFs were harvested and grown *in,vitro* before being chemically fixed and fluorescently labeled for three cytomorphological structures: the microtubule cytoskeleton (α-tubulin), nucleus (DAPI), and mitochondria (mtHsp70) (Fig. 1B). In numerous studies, morphological changes in these structures have been observed concomitant with changes in cellular function (*23-25*), suggesting potential links between cytomorphological and functional heterogeneity. Next, we developed image analysis algorithms to quantify 205 shape, size, and textural features (Table S3) in each cell, analyzing a 904-cell dataset. The distribution of cells in cytomorphological state space were visualized using Principal Component Analysis (PCA), a technique that converts high-dimensional datasets to lower-dimensional datasets by identifying linear combinations of features that represent the dataset with minimal loss of feature variance. These linear combinations are termed Principal Components (PCs) and are weighted for high-covariance features that together account for a large portion of population-wide variance. As discussed previously, many of our features exhibit high covariance (Fig. 1C); PCA allows for multiple co-varying features to be linearly combined into a single PC, thereby reducing the dimensionality of the dataset. In our dataset, PC1 was dominated by cell and nuclear morphometric features, while PC2 was dominated by cell and nuclear textural features, and PC3 dominated by nuclear textural and morphometric features (Table S4). Furthermore, plots of PC1, PC2, and PC3 (Fig, 1C, D), together accounting for 43.8% of total variance, revealed that WT MEFs occupy a set of cytomorphological states lying along a continuum, rather than discrete subsets of states.

**Fig. 1.**
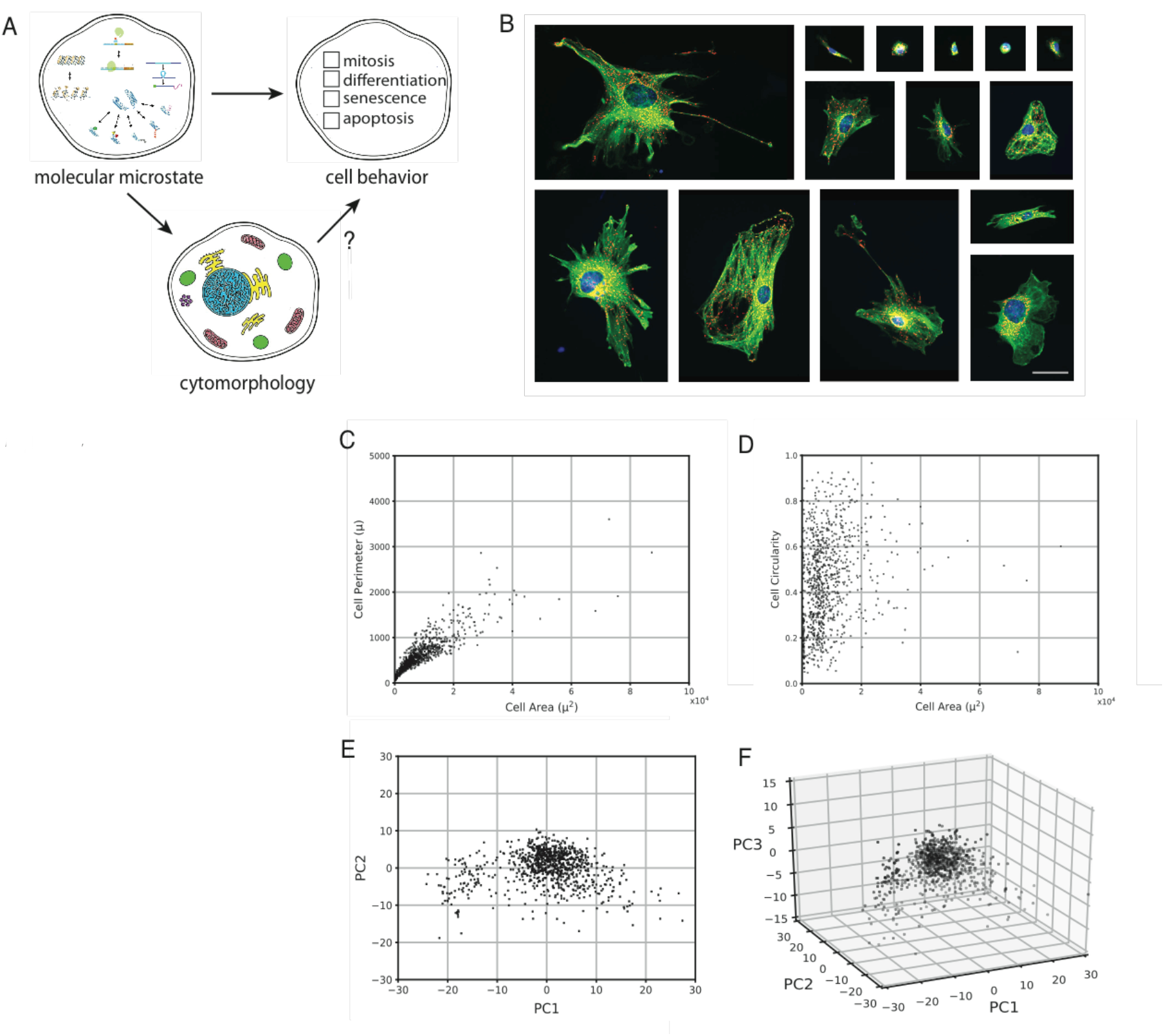
Morphological heterogeneity in WT MEFs. (**A**) Relationships between cell states at different levels of description. (**B**) Fluorescence images of fixed WT MEFs. The microtubule cytoskeleton (α-tubulin, green), mitochondria (mtHsp70, red), and nucleus (DAPI, blue) are labeled. Scale bar = 50μm. (**C** and **D**) Scatter plots of cell perimeter vs. cell area (**C**) and cell circularity vs. cell area (**D**) in a sample of fixed WT MEFs (n = 904). (**E** and **F**) Scatter plots of single-cell coordinates of fixed WT MEFs (n = 904) in PC1 vs. PC2 (**E**) and PC1 vs. PC2 vs. PC3 (**F**) space.

We next turned our attention to developing a dynamic map of transitions within state space by tracking cytomorphological changes in living cells. To achieve live cell imaging, we employed a lentivirus-based approach to introduce fluorescent reporters targeted to our structures of interest. Our lentivirus vector consisted of an EF1α promoter driving expression of fluorescent reporters localizing to the microtubule cytoskeleton (EGFP-tubulin), nucleus (mIFP-H2B), and mitochondria (tdTomato-mito-7) (Fig. 2A). Once transduced with this lentivirus, MEFs were imaged at 4h intervals over a 60h time course (Fig. 2B) and each cell’s location within state space calculated at each time point. New principal components (PCs) were calculated for living cells and a new cytomorphological state space constructed (Fig. S2); the topography of this space bears close resemblance to that observed for fixed cells (Fig. 1C). Next, cytomorphological state space transitions were plotted into a 60 × 60 bin representation of state space (Fig. 2C), where transition vectors were defined as originating at (PC1_*t*_, PC2_*t*_) and terminating at (PC1_*t+4h*_, PC2 _*t+4h*_). This plot revealed non-uniform magnitudes of transition, with vectors originating more centrally in state space of smaller magnitude than those originating peripherally. To quantify this observation, we plotted a heat map of the mean magnitude of transition of vectors originating from each bin (Fig. 2D) and confirmed that the magnitude of transition varies as a function of space state. In biological terms, this indicates that the degree of cytomorphological change within a given time interval varies as a function of the cytomorphological state of a cell.

**Fig. 2.**
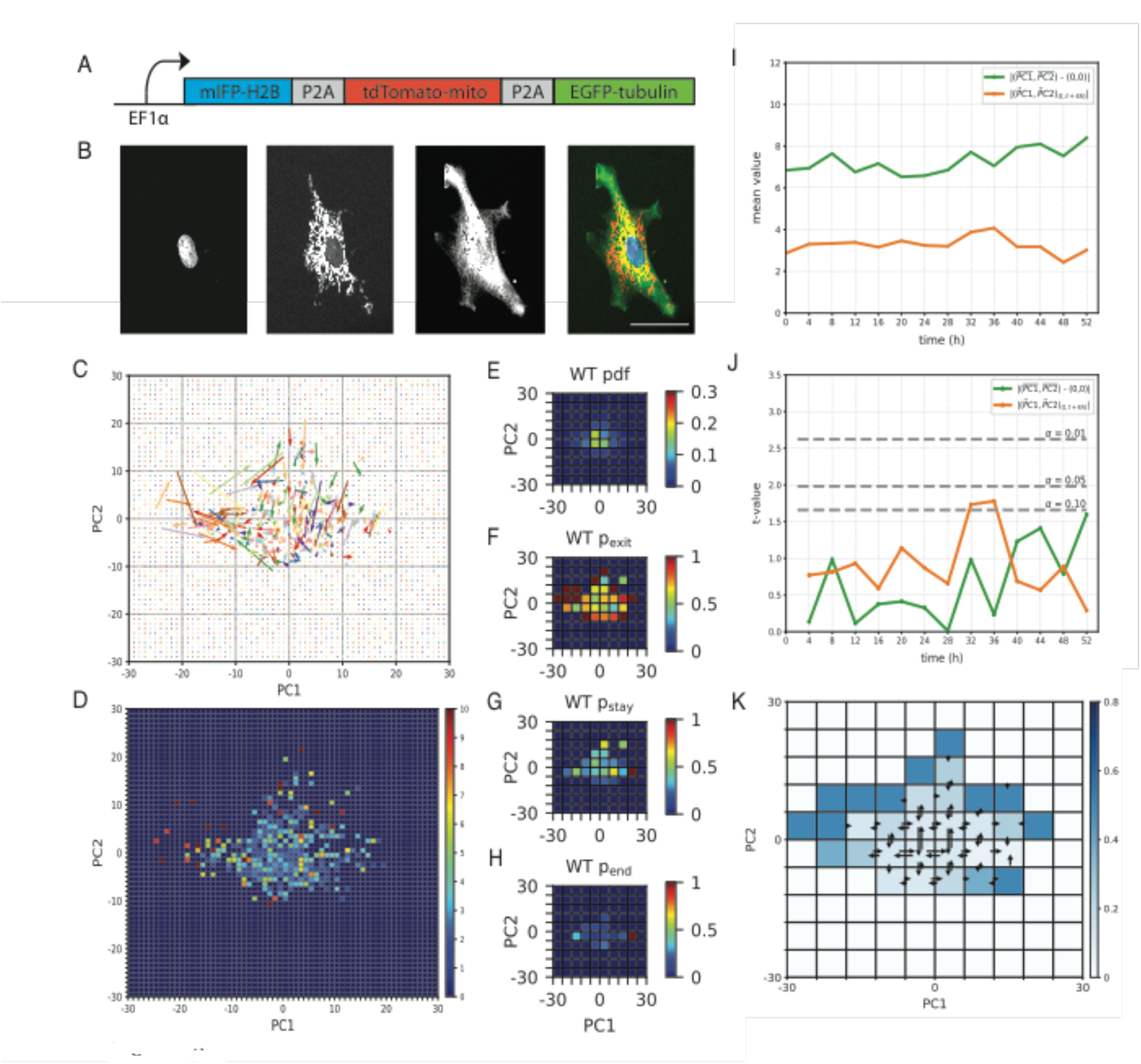
State space dynamics of living cells approximate a system at equilibrium. (**A**) Schematic of the fluorescent reporter lentivirus cassette. (**B**) Fluorescence images of a WT MEF expressing the lentivirus fluorescent reporter. From left to right: mIFP-H2B (nucleus), tdTomato-mito (mitochondria), EGFP-tubulin (microtubule cytoskeleton), composite. Scale bar = 50μm. (**C**) Experimentally observed transition vectors of WT MEFs over a 60h time course. Vectors originate at (PC1_t_, PC2_t_) and terminate at (PC1_t+4h_, PC2_t+4h_). (**D**) Heat map of the mean magnitude of transition vectors originating from each bin of a 60×60 bin representation of state space (**E through H**) Heat maps of the pdf (E), p_exit_ (F), p_stay_ (G), and p_end_ (H) of WT MEFs. (**I**) The distance from the origin of the mean (PC1, PC2) coordinate (green) and the mean transition vector magnitude from (PC1_t_, PC2_t_) to (PC1_t+4h_, PC2_t+4h_) (orange) of individual WT MEFs (n = 68). (**J**) Steady-state analysis. Two-sample t-test comparing, at each time point, relative to t = 0h, the distance from the origin of the mean (PC1, PC2) coordinate (green) and the mean transition vector magnitude from (PC1_t_, PC2_t_) to (PC1_t+4h_, PC2_t+4h_) (orange). Significance values are shown in dashed lines. (**K**) Detailed balance analysis. Heat map of the binomial statistics-based probability of the observed ratio of forward and reverse transitions occurring by random chance. Transitions between adjacent bins are shown; a full transition map can be found in the Supplemental Material (Fig. S3).

To further refine our dynamic map of cytomorphology space, we plotted the probability density function (pdf) of WT MEF state space occupancy (Fig. 2G) in a 10 × 10 bin representation of state space and identified a peak in the central (0, 0) region of this space. We then plotted the probability, over a 60h time course, that a cell occupying a bin would exit that bin by the next time point, (p_exit_) (Fig. 2H) and found that WT MEFs occupying more peripheral states had a higher probability of exit than those occupying more central states. The probability of staying within the same bin between consecutive time points (p_stay_), a measure of short-term occupancy, was also higher within these central bins (Fig. 2I). However, when we plotted the probability of a cell occupying the same bin as that occupied at the end of its time course (p_end_) (Fig. 2J), a rough measure of long-term occupancy, the mean probability fell by more than four-fold. These results suggest that WT MEF morphology space is energetically non-uniform and supports a directional bias towards the set of states surrounding (0, 0), but that entry into this space does not impose long-term occupancy. Interestingly, in PC space, the origin (0, 0) represents the mean value of contributing features, suggesting the possibility of a morphology-control mechanism acting to return cells to this morphological “mean.”

While these observations lay the groundwork for an empirical understanding of the dynamics of cytomorphology space, we wanted to investigate whether state space dynamics could be generalized into a framework describing the laws governing these transitions. In particular, we were interested in investigating the potential applicability of equilibrium dynamics formalisms and determining the accuracy with which the behavior of the system could be described using this approach.

Systems at equilibrium are, by definition, at steady state, where the distribution of state variables is invariant over all timescales. More rigorously, systems at equilibrium are also in detailed balance, where state transitions are accompanied by reciprocal transitions of equal magnitude in the reverse direction (*26*). Violations of detailed balance have been detected in some cellular processes (*27,28*), raising the question of the degree to which equilibrium concepts may apply in living systems.

To assess the steady state characteristics of cytomorphology space, we identified two state variables describing the occupancy and transition dynamics of MEFs within this state space. The first state variable, representing state space occupancy, measured distance of each cell from the origin (0, 0) in state space and calculated the mean distance at each time point (Fig. 2I, green). The second state variable, representing the dynamics of state space transitions, measured the mean magnitude of transition vectors at each time point (Fig. 2I, orange). At each time point, the mean distance from the origin as well as mean magnitude of transition of the population were compared to the mean distances and magnitudes observed at time t_1_ (0h) using a two-sample t-test. The results indicated that both the mean distance from the origin (0, 0) and mean magnitude of transition were statistically invariant (α = 0.05) at time scales ranging from 4h to 52h, (Fig. 2J), indicating that cytomorphological state space is at steady state at the temporal resolution of our experiment. To test for detailed balance, we tabulated the frequency of state transitions between each pair of bins within our 10×10 representation of state space (Fig. 2K, S3), where a binomial test (α = 0.05) revealed statistically insignificant variability in the rates of forward and reverse transitions between paired bins. The results of the steady state and detailed balance analyses indicated that the dynamical behavior of MEFs within cytomorphology space approximated those of a system at equilibrium.

To assess the applicability of an equilibrium dynamics approach to describing population-level transition dynamics of MEFs in cytomorphology space, we developed a framework based on an adaptation of Maxwell-Boltzmann statistics. Maxwell-Boltzmann statistics are an equilibrium statistical mechanics-based formalism for describing state occupancies of a system in terms of a single-valued state function that plays the role of a potential energy (*26,29*). According to this formalism, the relative occupancies of the different energy states of a system are a function of differences in the energy levels of different states as well as the temperature (T_b_) of the system. In classical statistical mechanics, the Maxwell-Boltzmann temperature (T_b_) refers to the thermodynamic temperature of the system as measured in Kelvins (K). This concept of temperature was adapted to our system by drawing parallels between particle velocity vectors and state space transition vectors (see Supplemental Material). This allowed us to calculate an effective temperature (T_WT_) of our system. The concept of energy state occupancies were similarly adapted by drawing parallels between particle occupancy of energetic states and cell occupancy of cytomorphological states, as measured by the probability density function (pdf, Fig. 2E). The combination of effective temperature and state space occupancy data were used to calculate the effective energy landscape underlying WT MEF cytomorphology space (Fig. 3B, see Supplemental Material for methods). This energy landscape was characterized by a global minimum surrounding (0, 0) and lacked any additional local maxima or minima. In an equilibrium system, the future behavior of the system can be predicted based on its current state, as the velocity field in state space is determined by the gradient of the potential energy landscape (Fig. 3A). To assess how predictive this effective energy landscape was of the experimentally observed transition vector field (Fig. 3C), we plotted the vector field predicted by the gradient of the energy landscape (Fig. 3D). By calculating the vector dot product of corresponding bins in the observed and predicted vector fields and then averaging the dot product across state space, we were able to quantify directional similarity between vector fields. Our results indicated that the experimentally observed transition vector field aligned more closely with the transition vector field predicted by our inferred effective energy landscape than with transition vector fields predicted from any of 100 scrambled iterations of the energy landscape, created by randomly distributing the original set of energy levels throughout state space (Fig. 3E, F, S4). Though cells are, unequivocally, non-equilibrium systems, we demonstrate here that at appropriate spatiotemporal scales, equilibrium dynamics formalisms can succeed in describing the behavior of an ensemble of non-equilibrium cells.

**Fig. 3.**
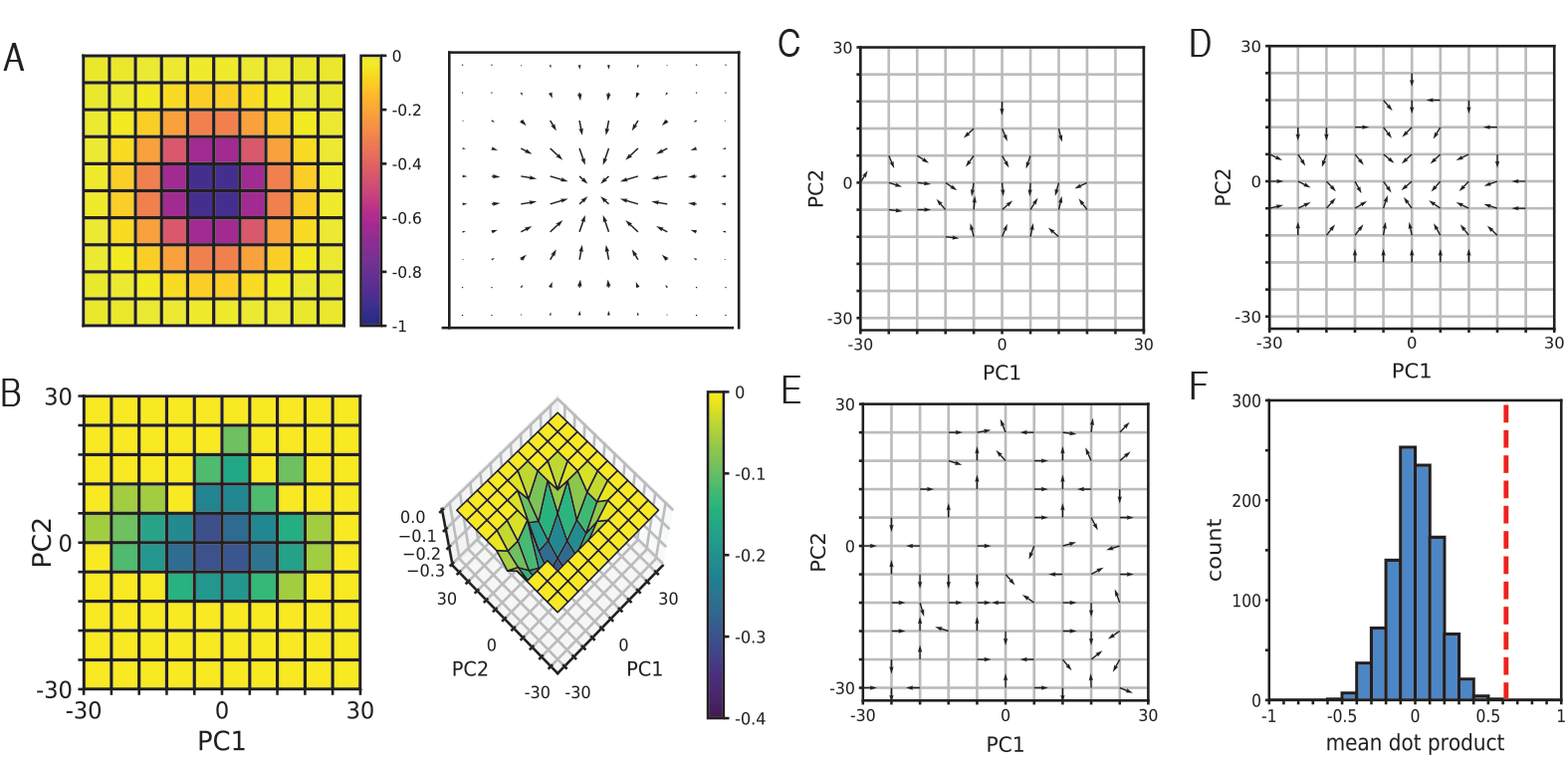
The gradient of an inferred effective energy landscape can predict experimentally observed transition dynamics. (A)Example of an energy landscape, formed by taking the inverse of a 2D Gaussian with μ_x_ = 0, μ_y_ = 0, σ_x_ = 1, and σ_y_ = 1(left) and the transition vector field (right) predicted by the gradient of that landscape. (B)Heat map (left) and surface plot (right) of the inferred effective energy landscape of WT MEF cytomorphology space. Energy values are scaled to a range of [−0.4, 0]. (**C** through **E**) State dynamics can be predicted from the gradient of the inferred effective energy landscape. Experimentally observed transition vectors (C), vectors derived from the gradient of the inferred effective energy landscape (D), and vectors derived from a scrambled energy landscape (E). Vectors are unit normalized. (**F**) Histogram of the mean dot product of the observed vectors (C) and vectors derived from 100 iterations of the scrambled landscape. The mean dot product of (C) and (D) is indicated by a red dashed line, indicating that the transition vector field predicted by the gradient of the inferred effective energy landscape is significantly predictive of experimentally observed transition dynamics.

Evidence shown here indicates that, within the context of specific parameter sets and spatiotemporal resolutions, the measured application of this approach can successfully describe the ensemble behavior of a population of non-equilibrium single cells. The application of equilibrium dynamics formalisms to describe a population of non-equilibrium systems does not imply equilibrium behavior at the single cell level and whether this same equilibrium formalism can be applied to individual cells is a separate question that we do not address here.

Having developed a framework for mapping cytomorphology space and a formalism for describing the energy landscape underlying this space, we were now in a position to investigate potential relationships between cytomorphological and functional variability. Given the many available examples of concomitant functional and cytomorphological changes, we hypothesized that the heterogeneity observed in cytomorphology space might correspond directly to functional heterogeneity within the population. To test this hypothesis, we screened five apoptosis drugs for conditions producing a heterogeneous response in a population of isogenic WT MEFs. We found that camptothecin (2μM), a topoisomerase I inhibitor, produced such a response, with a proportion of cells undergoing apoptosis within 96h of exposure, while others remained alive up to 21 days post-exposure (Fig. S5). The long duration over which some cells survived relative to the timescale of apoptosis suggests that the observed heterogeneity resulted from fundamentally different decisions in whether or not to undergo apoptosis, rather than from a simple delay in apoptosis.

Given many lines of evidence suggesting links between cytomorphology and cellular function, we investigated the possibility that intercellular variability in apoptotic response might be directly correlated with cytomorphological variability. To test this hypothesis, we transduced a population of MEFs with our cytomorphology-targeted fluorescent reporter lentivirus and imaged cells at 4h intervals, this time adding 2μM camptothecin at the 2h time point. Over the course of a 64h time course, we observed that 17 of 33 (52%) cells underwent apoptosis (Apoptosis[+]), while 16 of 33 (47%) cells did not (Apoptosis[−]).

We hypothesized that the response of a cell might correlate directly with its cytomorphological state prior to apoptosis induction, but when the space states of Apoptosis[+] and [−] cells at time 0h were plotted (Fig. S6) and analyzed, they were found to be statistically indistinguishable (Fig. S7). Though the initial space state was non-informative, an examination of the state transitions of each class of cell revealed characteristic differences in the dynamics of their state space transitions. The transition vectors of Apoptosis[+] cells (Fig. 4A, B) were of larger-than-average magnitude relative to the total cell population (Fig. S8) and displayed a directional bias towards the (−30, 30) region of PC space (Fig. S9). In contrast, the transition vectors of Apoptosis[−] cells (Fig. 4G, H) were of smaller-than-average magnitude (Fig. S8) and displayed a directional bias towards the (30, 0) region of PC space (Fig. S9). Interestingly, both groups of cells were also found to populate the central (0,0) region of PC space. Plots of the probabilities of exit from each bin (p_exit_) (Fig. 4D, J) revealed that both Apoptosis[+] and [−] cells exhibited higher probabilities of exit from peripheral states relative to central states. Plots of the probability of staying within the same bin during consecutive time points (p_stay_) (Fig. 4E, K), a measure of short-term occupancy, revealed a region of increased probability near (0, 0), in agreement with our model of an energetically favorable subspace surrounding this region. The p_stay_ of corresponding bins differed by no more than 1.5-fold between Apoptosis[+] and [−] cells, but plots of the probability of a cell occupying the same bin as that occupied at the end of its time course (p_end_) (Fig. 4F, L), a measure of long-term occupancy, revealed that long-term occupancy in Apoptosis[+] and [−] cells was restricted to different, non-overlapping regions within PC space. Apoptosis[+] cells exhibited increased p_end_ probabilities in the upper left (−30, 30) region of PC space, whereas Apoptosis[−] cells exhibited increased p_end_ probabilities in the central (0, 0) and far right (30, 0) regions of this space. These observations suggest that variability in the occupancy of Apoptosis [+] and [−] cells in WT MEF morphology space might contribute to functional divergence within the population. To statistically test this hypothesis, we ran two-dimensional Kolmogorov-Smirnov significance tests (α = 0.05) between pairs of corresponding probability plots (*30*) (Fig. 4M, S10), which revealed statistically significant differences between the p_exit_, p_stay_, and p_end_ plots of WT, Apoptosis[+] and Apoptosis[−] cells. These results suggest that camptothecin treatment produces statistically significant changes in the dynamics of WT MEF morphology space and that these changes lead to morphological and functional divergence within an isogenic population.

**Fig. 4.**
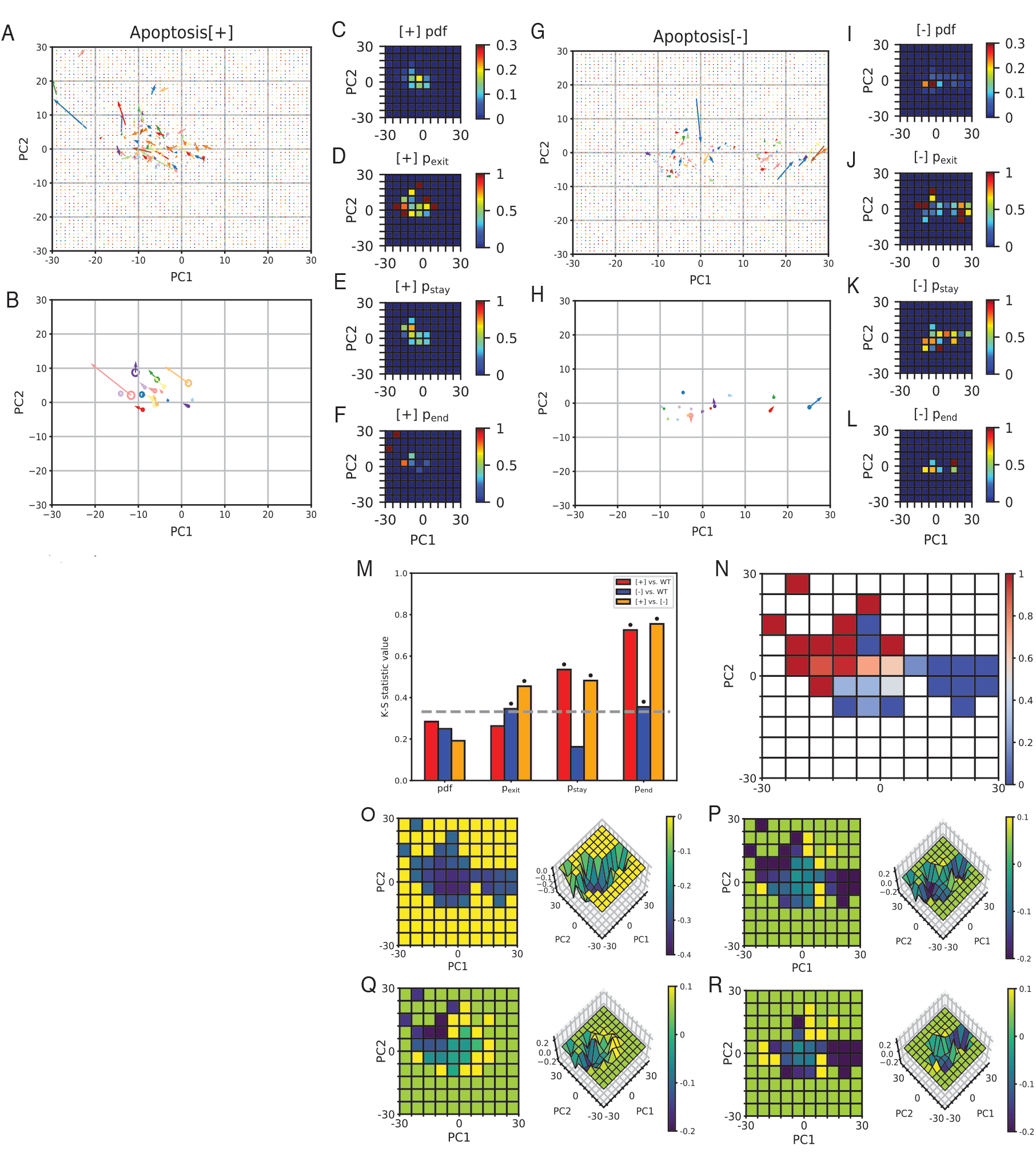
Morphology space state variability corresponds to variability in apoptosis response. (**A** and **G**) Experimentally observed transition vectors of Apoptosis[+] (A) and Apoptosis[−] (G) cells overs a 64h time course. Vectors originate at (PC1_t_, PC2_t_) and terminate at (PC1_t+4h_, PC2_t+4h_). (**B** and **H**) Single-cell mean transition vectors and mean (PC1, PC2) coordinates of Apoptosis[+] (B) and Apoptosis[−] (H) cells over a 64h time course. Radii of circles are scaled to the (PC1, PC2) coordinate variance over a 64h time course. (**C** through **F, I** through **L**) Heat maps of the pdf (C, I), p_exit_ (D, J), p_stay_ (E, K), and p_end_ (F, L) of Apoptosis[+] (C through F) and Apoptosis[−] (I through L) cells. (**M**) Bar plot of Kolmogorov-Smirnov significance values comparing the probability distributions of WT, Apoptosis[+], and Apoptosis[−] cells. The K-S significance value at α = 0.05 is indicated by a dashed line and statistically significant values are indicated by asterisks. (**N**) Heat map of the probability of apoptosis given occupancy of a particular state space bin at any point during the 64h time series. (**O**) Inferred effective energy landscape of camptothecin-treated WT MEFs. (**P** through **R**) Differences between the effective energy landscapes of WT and camptothecin-treated (P), Apoptosis[+] (Q), and Apoptosis[−] (R) cells.

To identify the underlying changes to cytomorphology space driving this behavior, we returned to our framework for calculating the effective energy landscape of a given state space. The transition vectors of camptothecin-treated cells were, on average, of 36% smaller magnitude than those of untreated cells (Fig. S11). In a system defined by Maxwell-Boltzmann statistics, smaller average displacements correspond to a lower effective temperature of the system. To test whether this “cooling” of morphology space could sufficiently explain observed changes in the probability density function (pdf) absent changes to the underlying energy landscape, we calculated the expected pdf of WT MEFs over a range of temperatures. We then ran a series of two-dimensional Kolmogorov-Smirnov (K-S) tests to quantify similarities between the observed and expected pdfs. A plot of K-S statistic values as a function of temperature (Fig. S12) revealed that similarities between the observed and expected pdfs were maximized at an effective temperature of 1.04T_WT_, suggesting that a change in temperature alone was insufficient to explain the observed change in the pdf and that camptothecin had fundamentally altered the underlying energy landscape. This new camptothecin-treated energy landscape was calculated (Fig. 4O) and the difference relative to the untreated energy landscaped plotted (Fig. 4P). This plot revealed that camptothecin treatment produced new energy minima in the upper left (−30, 30) and far right (30, 0) regions of PC space while further deepening the existing minimum at (0, 0).

Though both Apoptosis[+] and [−] cells effectively occupied the same camptothecin-treated energy landscape, we wanted to better understand whether specific features of this landscape explained their divergent behaviors. To do this, we separately calculated the effective temperatures of Apoptosis[+]and [−] cells and plotted their effective energy landscapes (Fig. S14, S15), along with their differences relative to the untreated energy landscape (Fig. 4Q, R). These plots revealed that divergent behaviors could be explained by spatially localized changes to the energy landscape. The effective energy landscape of Apoptosis[+] cells resembled that of untreated cells, with the exception of a new energy well in the upper left (−30, 30) region of PC space. This well coincided with the endpoint of 92% of Apoptosis[+] cells, indicating that this energy minimum acts as a “death” well into which apoptotic cells enter but cannot escape. In contrast, the energy landscape of Apoptosis[−] cells was characterized by a dramatic deepening in the far right (30, 0) region of PC space, accompanied by a second milder deepening in the central (0, 0) region. We hypothesized that these energy minima might act as protective barriers to apoptosis by hindering cell exit and minimizing the probability of cell entry into the (−30, 30) “death” well.

To test this possibility, we plotted the probability of apoptosis as a function of state space occupancy (Fig. 4N) and observed that entry at any point in the time series into states near (30, 0) corresponded to a 0% probability of apoptosis, whereas entry at any point in the time series into states in the (−30, 30) “death” well corresponded to a 100% probability of apoptosis. In between these two extremes were the central states surrounding (0, 0), where the probability of apoptosis was low, but nonzero. These findings support an interpretation where the addition of camptothecin produces three functionally distinct energy minima in WT MEF cytomorphology space: one acting as an irreversible “death” well into which cells enter and do not escape, and the other two acting as energetically favorable sub-regions that serve as protective barriers to apoptosis, one conferring absolute protection and the other conferring significant but incomplete protection.

We present here evidence demonstrating that heterogeneity in the topography and dynamics of cytomorphology space can effectively explain observed functional heterogeneity within an isogenic population. The discovery that drug exposure fundamentally alters the energetics of cytomorphology space lends insight into the effects of exogenous agents on cellular function at both the single cell and population scales. These observations further our understanding of the mechanisms driving functional heterogeneity and present a theoretical framework to guide future studies. These results strongly indicate that cytomorphological heterogeneity may be a functionally consequential mode of non-genetic heterogeneity, sister to the more widely studied forms of transcriptional and epigenetic heterogeneity.

As the basic functional unit of all living organisms, cells are continuously engaged in active biochemical processes and, from a microstate perspective, are distinct non-equilibrium systems. We demonstrate here that equilibrium-based formalisms can offer insight into the principles underlying non-genetic heterogeneity, suggesting that at coarse-grained scales, formalisms describing systems at equilibrium may be useful in describing and understanding the behaviors of non-equilibrium systems. If proven applicable in additional contexts, this principle may lead to the development of novel equilibrium statistical mechanics-based frameworks for studying an inherently non-equilibrium system, the cell. This approach has the potential to significantly further our understanding of biological systems, with recent work demonstrating promising applications of this approach (*31*). By illuminating the biophysical principles intertwining the molecular, morphological, and functional states of a cell, we hope to make progress in further uncovering the fundamental principles that govern all biological systems.

## Acknowledgements

We thank B. Panning and K. Leung for vector cloning guidance and use of tissue culture facilities, D. Ruggero, M. Truitt and M. McMahon for supply of MEFs, D. Larsen and S. Chen for microscopy support, and D. Yllanes and G. Huber for helpful discussion. This work was supported by an NSF Graduate Research Fellowship (A. C.), NSF grant MCB1515456 (W. M.), NIH grant GM097017 (W.M.), and the Center for Cellular Construction, NSF grant DBI1548297 (W. M.).

## List of Supplementary Materials

Materials and Methods

Supplementary Text

Figs. S1 to S15

Tables S1 to S5

Movies S1 to S3

