## Supplemental Material for "Dynamics of living cells in a cytomorphological state space"

**Materials and Methods**

1. Sample Preparation and Imaging

*Production of WT MEFs:* Primary mouse embryonic fibroblasts isolated from E13.5 embryos of C57BL/6 mice (Jackson Laboratory, #000664) were obtained from the lab of Davide Ruggero (UCSF Department of Urology). Passage 1 (P1) MEFs were flash frozen in cell culture freezing media (FBS + 10% DMSO) and stored at -80°C.

*Preparing chemically fixed WT MEFs for immunofluorescence imaging:* P1 WT MEFs were plated into 384-well tissue culture plates (Greiner, #781091) in full cell culture media (DME-H21 + 10% FBS + 1x pen/strep) at a density of 50 cells/well. Cells were incubated for 24h under 37°C, 5% $CO_2$ conditions, then chemically fixed and fluorescently labeled according to the following protocol:

1) Incubate cells in 4% paraformaldehyde (Electron Microscopy Sciences, #15710) for 15 minutes at room temperature.
2) Wash cells 1x with PBS.
3) Incubate cells in 0.1% Triton X-100 for 15 minutes at room temperature.
4) Wash cells 1x with PBS.
5) Incubate cells in blocking buffer (3% BSA + 5% normal goat serum (Jackson ImmunoResearch Labs, #005-000-121) + 0.1% Triton X-100) for 10 minutes at room temperature.
6) Incubate cells in 400x diluted mouse mAb mtHsp70 (ThermoFisher, #MA3-028) in blocking buffer for 1 hour at room temperature.
7) Wash cells 1x with PBS.
8) Incubate cells in 1000x diluted goat anti-mouse IgG AlexaFluor 488 (Life Technologies, #A11029) in blocking buffer for 1 hour at room temperature.
9) Wash cells 1x with PBS.
10) Incubate cells in 600x diluted rabbit pAb α-tubulin (abcam, #ab18261) in blocking buffer for 1 hour at room temperature.
11) Wash cells 1x with PBS.
12) Incubate cells in 1000x diluted goat anti-rabbit IgG AlexaFluor 647 (Life Technologies, #A21245) in blocking buffer for 1 hour at room temperature.
13) Wash cells 1x with PBS.
14) Incubate cells in 1μg/mL DAPI (Invitrogen, #D1306) in 0.1% Triton X-100 for 10 minutes at room temperature.
15) Wash cells 3x with PBS. Store cells at 4°C in PBS, protected from light, for up to 3 days prior to imaging.

*Fluorescence imaging of chemically fixed WT MEFS:* WT MEFs in 384-well plates were imaged on an IN Cell Analyzer 2000 (GE Healthcare, #28-9798-74) fitted with a Nikon Plan Fluor 40x (NA 0.60) objective. Cells were imaged in the DAPI, FITC, and Cy5 channels to illuminate the nucleus, mitochondria, and microtubule cytoskeleton, respectively.

*Construction of fluorescent reporter lentivirus expression vector:* For the vector backbone, we used the pWPXL vector (Addgene, #12257), a lentivirus expression vector containing a GFP reporter driven by an EF1α promoter. We excised the GFP reporter and replaced it with a cassette containing, in tandem, fluorescent reporters localizing to the microtubule cytoskeleton (EGFP-α-tubulin), mitochondria (tdTomato-mito-7), and nucleus (mIFP-H2B). These reporters were separated by copies of P2A (66bp) (*32*), a self-cleaving peptide that allowed for production of three separate fluorescent reporters from a single transcript. Cloning of the vector was accomplished in three steps:

1) Fluorescent reporter plasmids for our three structures of interest were acquired from the Davidson Collection stored at the Nikon Imaging Center (NIC) at UCSF. The plasmids were: *EGFP-tubulin-6* (α-tubulin linked to an enhanced GFP), *mIFP-H2B-6* (histone H2B linked to a monomeric infrared fluorescent protein), and *tdTomato-mito-7* (a mitochondria localization sequence linked to a tdTomato fluorescent protein). Reporters were separated by copies of P2A (66bp), resulting in a single cassette with EF1α driving expression of all three fluorescent reporters. Cloning was accomplished using a Gibson Assembly approach. Primers were purchased from Integrated DNA Technologies (IDT); sequences are listed in (Table S1).

a) pWPXL homology was added to the 5' end of mIFP-H2B-6 using the F-H2B_KOZAK primer; this primer preserves the BamHI restriction site and introduces a Kozak sequence to increase translation efficiency.

b) The P2A sequence (66bp) (*32*) was added to the 3' end of mIFP-H2B-6 using the R-mIFP_EXTENSION primer, forming mIFP-H2B-6-P2A.

c) The tdTomato-mito-7 construct was separated into two modules: tdTomato1, with 5' P2A homology and 3' tandem linker homology, and tdTomato2, with 5' tandem linker homology and 3' P2A homology. It was necessary to PCR amplify the two modules separately because the tandem tdTomato modules are homologous, resulting in random primer attachment to both modules if PCR amplified as a single construct. P2A homology was added to the 5' end of tdTomato1 using the F-MITO primer. Tandem linker homology was maintained at the 3' end of tdTomato1 using the R-tdTomatoLinker primer. Tandem linker homology was maintained at the 5' end of tdTomato2 using the F-tdTomatoLinker primer. P2A-EGFP-tubulin-6 homology was added to the 3' end of tdTomato2 using the R-MITO primer.

d) The P2A sequence (66bp) (*32*) was added to the 5' end of EGFP-tubulin-6 using the F-EGFP_EXTENSION primer.

e) pWPXL homology was added to the 3' end of EGFP-tubulin-6 using the R-TUBULIN primer.

2) Primers were designed to produce fragments compatible with Gibson Assembly, with 16-42bp homology between fragments. Fragments were PCR amplified using Phusion High Fidelity DNA Polymerase (NEB, #M0530L) following a standard PCR protocol. Fragments were stitched together using a Gibson Assembly Kit (NEB, #E2611S) following kit instructions. PCR amplification and Gibson Assembly were accomplished in the following order:

a) PCR amplification of mIFP-H2B-6 to obtain mIFP-H2B-6-P2A, with 5' pWPXL homology and a 3' P2A sequence.

b) PCR amplification of tdTomato2-mito-7 to obtain tdTomato1, the first tdTomato module, with 5' homology to P2A and 3' homology to the tandem linker.

c) PCR amplification of tdTomato2-mito-7 to obtain tdTomato2, the second tdTomato module, with 5' homology to the tandem linker and 3' homology to P2A.

d) PCR amplification of EGFP-tubulin-6 to obtain P2A-EGFP-tubulin-6, with a 5' P2A sequence and 3' pWPXL homology.

e) Gibson Assembly of mIFP-H2B-6-P2A and tdTomato1 to obtain mIFP-H2B-6-P2A-tdTomato1, with 5' pWPXL homology and 3' tandem linker homology.

f) Gibson Assembly of tdTomato2 and P2A-EGFP-tubulin-6 to obtain tdTomato2-P2A-EGFP-tubulin-6, with 5' tandem linker homology and 3' pWPXL homology.

g) PCR amplification of mIFP-H2B-6-P2A-tdTomato1 using primers F-H2B_KOZAK and R-tdTomatoLinker, and PCR amplification of tdTomato2-P2A-EGFP-tubulin-6 using primers F-tdTomatoLinker and R-TUBULIN.

h) Full Gibson Assembly of mIFP-H2B-6-P2A-tdTomato1 and tdTomato2-P2A-EGFP-tubulin-6 to obtain the full fluorescent reporter cassette.

3) The assembled product was transformed into 5α Competent *E.coli* cells (NEB, #C2987) following kit instructions. Cells were plated onto appropriate antibiotic- selection LB agar plates and incubated at 37°C overnight. Colonies were screened using a standard colony PCR protocol and hits cultured and sent for full-coverage sequencing of the complete fluorescent reporter cassette. Successful hits were transformed into 293T cells to ascertain expected fluorescence and cytostructural localization. A single hit was then aliquoted and stored at -20°C until ready for use.

*Producing fluorescent reporter lentivirus:* A standard lentivirus production protocol was used to produce lentivirus in 293T cells. Lentivirus product was concentrated with Lenti-X Concentrator (Clontech, #PT4421-2) and titered in WT MEFs to identify conditions producing maximum transduction (as determined by fluorescence) with minimal cell death. Lentivirus was then stored in single-use aliquots at -80°C until ready for use.

*Preparing fluorescent reporter-expressing WT MEFs for live-cell imaging:* P1 WT MEFs were seeded into 96-well tissue culture plates (Greiner, #655090) in antibiotic-free cell culture media (DME H-21 + 10% FBS) at a density of 5000 cells/well. Cells were incubated at 37°C, 5% $CO_2$ for 24 hours and the appropriate titer of fluorescent reporter lentivirus added. After an additional 24 hours, cells were trypsinized and re-seeded into a new 96-well tissue culture plate (Greiner, #655090) in full cell culture media (DME H-21 + 10% FBS + 1x pen/strep) at a density of 500 cells/well. After an additional 24 hours, media was replaced with full cell culture media containing 100μM biliverdin hydrochloride (BV; Sigma, #30891). BV is a cofactor to mIFP and integral to mIFP fluorescence. After an additional 24 hours, BV-containing media was removed and cells washed 3x with PBS and re-incubated under full cell culture media (DME H-21 + 10% FBS + 1x pen/strep).

*Imaging of fluorescent reporter-expressing WT MEFs:* Cells were imaged at the Nikon Imaging Center (NIC) at UCSF on a spinning disk confocal microscope (Nikon Ti inverted fluorescence microscope fitted with an Andor Borealis CSU-W1 confocal). The microscope was fitted with a Plan Apo λ 40x objective (NA 0.95) and equipped with 405, 488, 561, and 640nm laser lines and 450/50m (DAPI), 525/50m (FITC), 600/50m (Cy3), and 700/75m (Cy5) filters. A stage-top temperature and $CO_2$ control chamber was set to 37°C and 0.3 l/min to mimic tissue culture conditions. Cells were imaged in the FITC, Cy3, and Cy5 channels at 2h intervals over a 60h time course.

*Screening apoptosis drugs and concentrations:* For our studies of functional heterogeneity in WT MEFs, we chose to investigate apoptosis as our functional readout. We screened a panel of five apoptosis inducers (abcam, #ab102480) across a range of concentrations (Table S2) and identified camptothecin (2μM) as the most promising candidate, as it only produced apoptosis in a subset of cells, indicating a heterogeneous response.

*Imaging of fluorescent reporter-expressing WT MEFs under apoptosis conditions:* Cells were prepared as detailed above ("*Preparing fluorescent reporter-expressing WT MEFs for live-cell imaging*"). Cells were imaged using the above mentioned microscopy set-up ("*Imaging of fluorescent reporter-expressing WT MEF*"). Cells were imaged at 2 hour intervals over a 64 hour time course, with 2μM camptothecin added after the second (2 hour) time point. This experimental set-up provided cytomorphological information both pre- and post-camptothecin addition.

2. Image Analysis and Feature Extraction

*Preparing images for analysis:* To achieve as homogeneous a population as possible, only single, isolated cells that were not in contact with other cells nor with the well edge were chosen for analysis. Furthermore, only cells that were visually determined to be in the G1, S, or G2 phases (not in M phase) were chosen for analysis. In the fixed cell experiments, cells occasionally spanned more than one field of view (FOV). To include these cells in downstream analysis, adjacent FOVs were stitched together using the ImageJ "Pairwise Stitching" plug-in (*33*). In the live-cell experiments, cells that were not captured within a single FOV were not analyzed.

*Image segmentation:* Cells were first segmented on the FITC (EGFP-tubulin) channel to identify a cell outline. Segmentation was accomplished using a custom segmentation algorithm written in MATLAB. This algorithm consisted of a series of dilation, filling, erosion, and smoothing steps. Once the cell outline was established, pixels interior to the cell outline were further segmented on the Cy3 (tdTomato-mito) and Cy5 (mIFP-H2B) channels. This was accomplished using a combination of pre-existing (*34*) and custom segmentation algorithms written in MATLAB that similarly consisted of a series of dilation, filling, erosion, and smoothing steps. These segmentation steps produced object masks for the microtubule cytoskeleton, nucleus, and mitochondria of each cell, from which morphometric and textural features were then extracted. A sample segmentation can be found in (Fig. S1). The custom segmentation and feature analysis scripts are available at https://github.com/amyrchang/WTMEF.

*Feature extraction:* A complete list of the 205 features along with their descriptions can be found in (Table S3).

a) *Microtubule cytoskeleton and nucleus:* The microtubule cytoskeleton and nucleus were both treated as single objects and analyzed for single-object features. Custom image analysis algorithms were written in MATLAB to extract cell and nucleus morphometric measurements (Table S3, Features 1-33; 34-72). These include features such as area, perimeter, and ellipticity, as well as more complex features such as the area of a convex hull surrounding the object and the proportional change in the object's area and perimeter upon smoothing of the perimeter. A second set of image analysis algorithms were designed to extract textural features from each object (Table S3, Features 73-104; 105- 136). These include explicit textural features such as contrast and homogeneity, as well as custom spatial distribution features that slice objects into concentric rings of increasing radius and analyze the distribution of features within each ring.

b) *Mitochondria:* In contrast to the microtubule cytoskeleton and nucleus, the mitochondrial object is composed of many individual, disconnected objects. We therefore designed our mitochondrial feature extraction algorithms to extract features of both individual mitochondrial objects as well as the sum of the objects. The first set of algorithms extracted morphometric features such as the area and perimeter both of individual mitochondrial objects and their sum object (Table S3, Features 137-173), as well as features measuring the connectivity of objects (e.g. number of nodes, ratio of fused vs. fragmented mitochondria, etc.). A second set of algorithms extracted textural and spatial intensity distribution features of the mitochondria (Table S3, Features 174- 205). These included measurements of the homogeneity of mitochondria distribution within the cell (e.g. are they uniformly distributed throughout the cell, or are they clustered?), as well as measurements of their spatial distribution within the cell (e.g. does there appear to be spatial bias towards either perinuclear or peripheral locations?)

### 3. Statistical Analysis

*Principal Component Analysis:* To dimensionally reduce the 205-feature space into a more interpretable, lower dimensional space, we employed principal component analysis (PCA). PCA works by weighting features according to their contribution to the overall intercellular variance of each feature and creating a linear combination based on these weightings, effectively forming a new, variance-weighted basis composed of Principal Components. We began by compiling each dataset into a matrix $M_{i,j}$ with 205 rows indexed by the letter *i* and 904 columns indexed by the letter *j*. Each column represented a different cell and each row a different feature. We then normalized the feature values within each column by calculating the μ (mean) and σ (std) of each feature and calculating the associated z-score of each measurement, producing $Z_{i,j}$, a matrix of z-scores, where $z_{(i,j)}$ = the z-value for the $i^{th}$ feature of the $j^{th}$ cell:

$$Z = \begin{bmatrix} z_{(1,1)} & z_{(1,2)} & \cdots & z_{(1,903)} & z_{(1,904)} \\ z_{(2,1)}, & z_{(2,2)} & \cdots & z_{(2,903)} & z_{(2,904)} \\ \cdots & \cdots & \cdots & \cdots & \cdots \\ z_{(204,1)} & z_{(204,2)} & \cdots & z_{(204,903)} & z_{(204,904)} \\ z_{(205,1)} & z_{(205,2)} & \cdots & z_{(205,903)} & z_{(205,904)} \end{bmatrix} \qquad Z^T = \begin{bmatrix} z_{(1,1)} & z_{(2,1)} & \cdots & z_{(204,1)} & z_{(205,1)} \\ z_{(1,2)} & z_{(2,2)} & \cdots & z_{(204,2)} & z_{(205,2)} \\ \cdots & \cdots\cdots & \cdots & \cdots & \cdots \\ z_{(1,903)} & z_{(2,903)} & \cdots & z_{(204,903)} & z_{(205.903)} \\ z_{(1,904)} & z_{(2,904)} & \cdots & z_{(204,904)} & z_{(205.904)} \end{bmatrix}$$

PCA requires a calculation of the covariance of features (Eq. 1). We eliminate the fraction in front of $ZZ^T$, as this is essentially a weighting factor and does not affect the calculation of eigenvectors or their weightings relative to each other. The values for the three eigenvectors with the largest eigenvalues, representing Principal Components (PC) 1, 2, and 3, are listed in (Table S4, S5). Separate PCAs were performed for the fixed and live cell datasets. $M_{i,j}$ matrices for fixed, live, [+], and [-] WT MEFs are available at https://github.com/amyrchang/WTMEF.

$$S_Z = \text{cov}(Z) = \frac{1}{904-1} ZZ^T \qquad \text{(Eq. 1)}$$

*Probability Plots (pdf, $p_{enter}$, $p_{end}$, $p_{stay}$, $p_{end}$):* Probability plot calculations were based on the cumulative behavior of all cells over the course of the time series, normalized by the number of times each cell was observed. Probability plots for each cell were calculated by summing the number of occurrences of a specific behavior within each bin and dividing by the total number of occurrences of that cell within each bin (Eq. 2-5). The mean probability for each behavior in each bin was then calculated across all probability plots to produce an average probability plot representing the behavior of all cells, with the behavior of each cell weighted equally in order to account equally for the behavior of each cell.

$$\text{pdf}_{(i,j)} = \frac{\left(\sum_n^N \frac{\#\ of\ occurrences\ of\ cell_n\ in\ bin_{(i,j)}}{total\ \#\ of\ occurrences\ of\ cell_n}\right)}{N} \qquad \text{(Eq. 2)}$$

$$p_{exit(i,j)} = \frac{\left(\sum_n^N \frac{\#\ of\ occurrences\ of\ cell_n\ exiting\ from\ bin_{(i,j)}}{total\ \#\ of\ occurrences\ of\ cell_n\ in\ bin_{(i,j)}}\right)}{N} \qquad \text{(Eq. 3)}$$

$$p_{stay(i,j)} = \frac{\left(\sum_n^N \frac{\#\ of\ occurrences\ of\ cell_n\ staying\ in\ bin_{(i,j)}\ in\ consecutive\ time\ points}{total\ \#\ of\ occurrences\ of\ cell_n\ in\ bin_{(i,j)}}\right)}{N} \qquad \text{(Eq. 4)}$$

$$p_{end(i,j)} = \frac{\left(\sum_n^N \frac{\#\ of\ occurrences\ of\ cell_n\ in\ bin_{(i,j)}\ ending\ the\ time\ course\ in\ bin_{(i,j)}}{total\ \#\ of\ occurrences\ of\ cell_n\ in\ bin_{(i,j)}}\right)}{N} \qquad \text{(Eq. 5)}$$

*Stationarity Analysis:* Two-sample t-tests were run to determine the statistical significance of changes in the population mean PC1 coordinate, mean PC2 coordinate, and mean vector magnitude during consecutive time points. There were 46 degrees of freedom (dof) for each of the datasets, with significance values of $t = 1.679$, $t = 2.013$, and $t = 2.687$ at significance levels of $\alpha = 0.10$, $\alpha = 0.05$, and $\alpha = 0.01$, respectively.

*Detailed Balance Analysis:* In our detailed balance analysis, we focused only on tracking reciprocal transitions between neighboring bins as opposed to transitions between all possible bins to render the analysis more manageable. We plotted arrows with magnitudes scaled to the number of transitions between neighboring bins and calculated the binomial probability of the observed ratio of forward to reverse transitions occurring by random chance. For the heat map, the lowest binomial probability for each bin was plotted (a bin may have more than one calculated binomial probability due to transitions in four possible directions away from the bin). The lowest binomial probability was 0.0625, which was higher than the significance value of $\alpha = 0.05$, indicating that even the most imbalanced reciprocal transition ratio had a greater than 5% probability of occurring by random chance.

*Effective Energy Landscapes:* We developed an adaptation of Maxwell-Boltzmann statistics, which describes the probability density function of a system as a function of the temperature and energy states of that system, and applied this to our two-dimensional Principal Component-based morphology space. We then used this adaptation to map out the inferred effective energy landscape underlying WT MEF morphology space.

The equation governing Maxwell-Boltzmann statistics is shown in (Eq. 6), where $\bar{n}_s$ is the average number of particles found in state *s* with energy level, $\xi_s$, *N* is the total number of particles in the system, *r* is the total number of energy states of the system, and $\beta = 1 / k_B T$, where $k_B$ is the Boltzmann constant ($k_B = 1.381 * 10^{-23}$ m$^2$ kg s$^{-2}$ K$^{-1}$) and *T* is the temperature of the system. *Z* represents the partition function, or the sum of $e^{-\beta\xi_s}$ over all possible energy states.

$$\frac{\overline{n_s}}{N} = \frac{e^{-\beta\xi_s}}{\sum_r e^{-\beta\xi_r}} = \frac{e^{-\beta\xi_s}}{Z} \qquad \text{(Eq. 6)}$$

We then used our experimental data to calculate values for several of the above variables. The value $\overline{n_s}/N$, is simply the probability density at each bin, while T is the "temperature" of WT MEF morphology space. To calculate an effective temperature for a given cell morphology space, we abstracted transition vectors to represent particle velocity vectors and used the equipartition theorem to derive temperature from the average velocity of a particle in a two-dimensional plane, where each available dimension of movement contributes $\frac{1}{2} * k_B T$ kinetic energy to the system. Given a two-dimensional plane, the total kinetic energy of the system is equal to $k_B T$. We then derived the following:

$$\frac{1}{2} m\bar{v}^2 = k_b T \qquad \text{(Eq. 7)}$$

$$T = \frac{m\bar{v}^2}{2k_b} \qquad \text{(Eq. 8)}$$

Using this approach, we calculated the average transition vector magnitude and, using an arbitrary mass of $10^{-21}$ kg/particle for unit conversion purposes, calculated an effective temperature, $T_{WT}$, of WT MEF morphology space. Because of the arbitrary mass assumption, the value of the effective temperature is not in itself meaningful; however, (Eq. 8) allows for the effective temperature to be calculated under different experimental conditions, allowing for experiments where cells show different overall rates of state transitions to be compared.

Without enough information to solve for the partition function, Z, we could not calculate absolute values for the energy states $\xi_1, \xi_2, \xi_3, \xi_4, \dots, \xi_s$ of the system. However, in the context of our energy landscape, we were primarily interested in the values of energy states *relative* to each other rather than their absolute values. By setting the partition function equal to $e^{-\beta\xi_0}$, where $\xi_0$ is an arbitrarily chosen reference value, (Eq. 6) can be rewritten as:

$$\frac{\bar{n}_x}{N} = \frac{e^{-\beta\xi_s}}{e^{-\beta\xi_0}} = e^{-\beta(\xi_s - \xi_0)} \qquad \text{(Eq. 9)}$$

A rearrangement of (Eq. 9) gives:

$$\xi_s - \xi_0 = \frac{\ln\left(\frac{\bar{n}_s}{N}\right)}{-\beta} \qquad \text{(Eq. 10)}$$

We set the reference value, $\xi_0$ ,equal to $0.4*10^{-19}$ J and used a Boltzmann constant of $k_b = 1.381*10^{-23}$ $m^2$ kg $s^{-2}$ $K^{-1}$, along with the WT probability density function and the calculated value of the effective WT temperature, $T_{WT}$, to calculate values for $\xi_1 - \xi_0$, $\xi_2 - \xi_0$, $\xi_3 - \xi_0$, … , $\xi_s - \xi_0$. To plot the energy values in easily interpretable units, we linearly modified these values by plotting the quotient of this value divided by $10^{-19}$.

*Calculating the Predicted Transition Vector Field:* In order to calculate the transition vector field predicted by our WT energy landscape, Q, we calculated the gradient of the landscape (Eq. 11), where *a* and *b* are the PC1 and PC2 vectors, respectively.

$$\nabla = \frac{\partial Q}{\partial a}\hat{a} + \frac{\partial Q}{\partial b}\hat{b} \qquad \text{(Eq. 11)}$$

*Scrambled Energy Landscapes:* To create a scrambled version of the energy landscape, we took the energy values calculated for the effective energy landscape and randomly assigned them to new bins within a 10x10 bin space. A sample scrambled energy landscape is shown in (Fig. S4). The transition vector field predicted by the scrambled energy landscape (Fig. S4) was similarly calculated by taking the gradient of the landscape according to (Eq. 11).

*Vector Field Dot Product Calculations:* To quantify the degree of similarity between vector fields, we unit normalized each vector field so that only directional information would factor into the dot product calculation. We then calculated the dot product for all bins for which there were vectors in both of the vector fields being compared. The mean value of these dot products was calculated and taken as an indication of the degree of directional similarity between vector fields. The data for the histogram in (Fig. 2P) was obtained by calculating 100 scrambled versions of the effective energy landscape (example found in Fig. S4), calculating their predicted transition vector fields (example found in Fig. 2O), and taking the mean dot product of each vector field with that observed experimentally. The value for the red asterisk in (Fig. 2P) was obtained by taking the mean dot product of the transition vector field predicted by the inferred effective energy landscape (Fig. 2N) with that observed experimentally (Fig. 2M).

*Comparison of Camptothecin Observed and WT Predicted Probability Density Functions:* To determine whether a change in effective temperature alone was sufficient to explain the observed change to the WT probability density function (pdf) upon camptothecin addition, we used our adaptation of Maxwell-Boltzmann statistics to predict the pdf of cells occupying the WT effective energy landscape at temperatures ranging from $T_{WT}\log_{10}x$, with x ranging from 1 to 50. We then ran two-dimensional Kolmogorov-Smirnov (K-S) tests to quantify the degree of similarity between each predicted pdf and that observed experimentally. A plot of K-S statistic values as a function of effective temperature (Fig. S12) indicated that the pdf predicted at an effective temperature of 1.04 $T_{WT}$ was most similar to that observed experimentally. However, the experimentally observed effective temperature of camptothecin morphology space was $0.41T_{WT}$, indicating that camptothecin treatment had altered the energy landscape underlying WT MEF morphology space.

*Camptothecin Effective Energy Landscape:* The camptothecin effective energy landscape was calculated using the same method used to calculate the WT effective energy landscape ("*Application of Maxwell-Boltzmann Statistics to Calculate Effective Energy Landscapes*").

*Calculating Changes to the Effective Energy Landscape:* Cells treated with camptothecin occupied space states that were unoccupied under untreated conditions, so a modification of the WT energy landscape to include energy level values for all bins was necessary to calculate differences across all bins. We modified the effective energy landscape of untreated WT MEFs by populating the bins for which no cells had been observed with the lowest probability density of an occupied bin ($p = 0.0017$). Since the true probability density of bins for which no cells were observed can be considered to be somewhere between 0 and 0.0017, populating these bins with the lowest probability density represented the "most challenging" scenario under which to identify a difference in the energy landscape. We then calculated a modified WT energy landscape (Fig. S13) using our Maxwell-Boltzmann adaptation and plotted the differences between the WT and camptothecin (Apoptosis[+] and Apoptosis[-] combined) (Fig. 3R), Apoptosis[+] (Fig. S14), and Apoptosis[-] (Fig. S15) energy landscapes.

*Identifying changes to the probability heat maps:* Two-dimensional Kolmogorov-Smirnov (K-S) tests were run on pairs of corresponding probability plots of untreated WT, Apoptosis[+], and Apoptosis[-] cells to determine whether there were statistically significant differences between corresponding probability plots. The K-S significance values were plotted and compared to a K-S significance value of 0.1391 for $\alpha = 0.05$ (Fig. 3O).

*Heat Map of the Probability of Apoptosis:* Calculations of the probability of apoptosis were based on cumulative bin occupancies and apoptotic response over the entire time course. The number of occupancies in each bin over the entire time course was counted (e.g, a single cell occupying the same bin over two time points counted as two occupancies). The number of occupancies of Apoptosis[+] cells in each bin over the entire time course was also counted. For each bin, the quotient of the second value divided by the first value represented the probability of apoptosis for a cell chosen at random to occupy that bin (Eq. 12). This probability was then calculated for all bins and plotted as a heat map (Fig. 3P).

$$p_{\text{apoptosis},(i,j)} = \frac{\#\ of\ occupancies\ of\ bin_{(i,j)}\ by\ an\ Apoptosis[+]\ cell}{total\ \#\ of\ occupancies\ of\ bin_{(i,j)}} \qquad \text{(Eq. 12)}$$

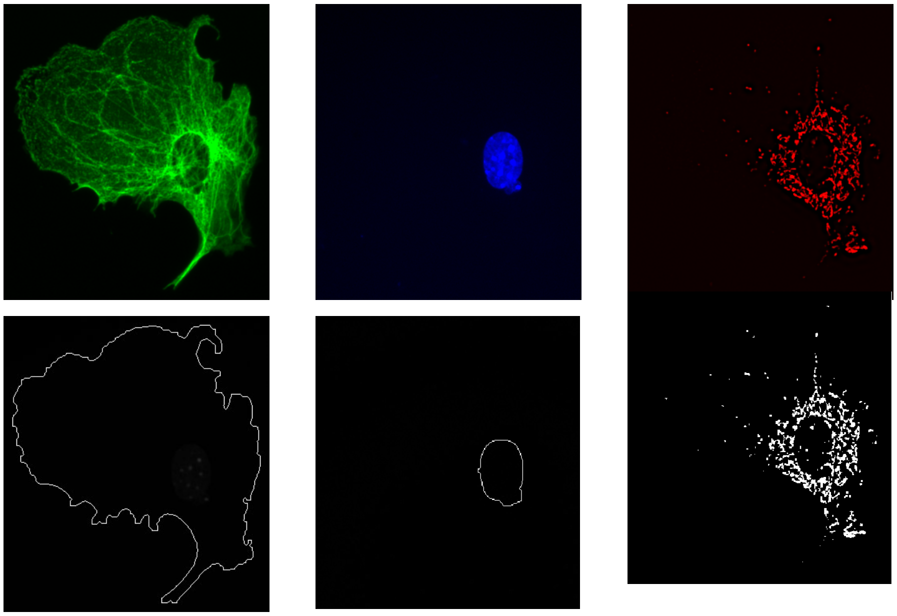

**Fig. S1. Segmentation of cytomorphological structures.** WT MEF fixed and fluorescently labeled for α-tubulin (top, left), DAPI (top, center), and mtHsp70 (top, right). Each structure was segmented using custom image segmentation algorithms written in MATLAB. The original image (top, colored) and segmented outline/mask (bottom, white) are shown for the microtubule cytoskeleton (left, green), nucleus (center, blue), and mitochondria (right, red). Image taken at 40x magnification.

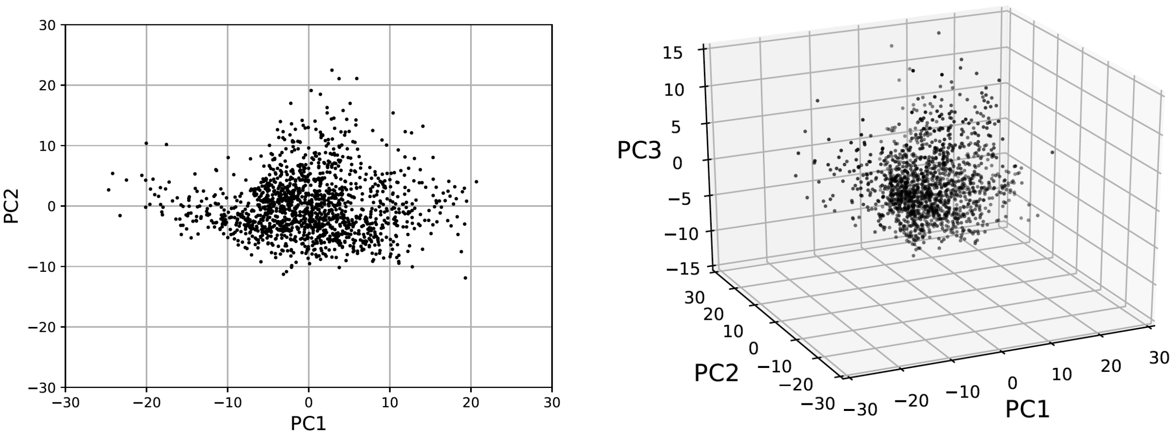


**Fig. S2. Distribution of Live WT MEFs in Principal Component Space.** Scatter plots of single-cell coordinates of live WT MEFs (n = 1432) in live cell PC1 vs. PC2 (left) and PC1 vs. PC2 vs. PC3 (right) space. PC1, PC2, and PC3 account for 23.4%, 11.8%, and 8.6%, of the total variance, respectively.

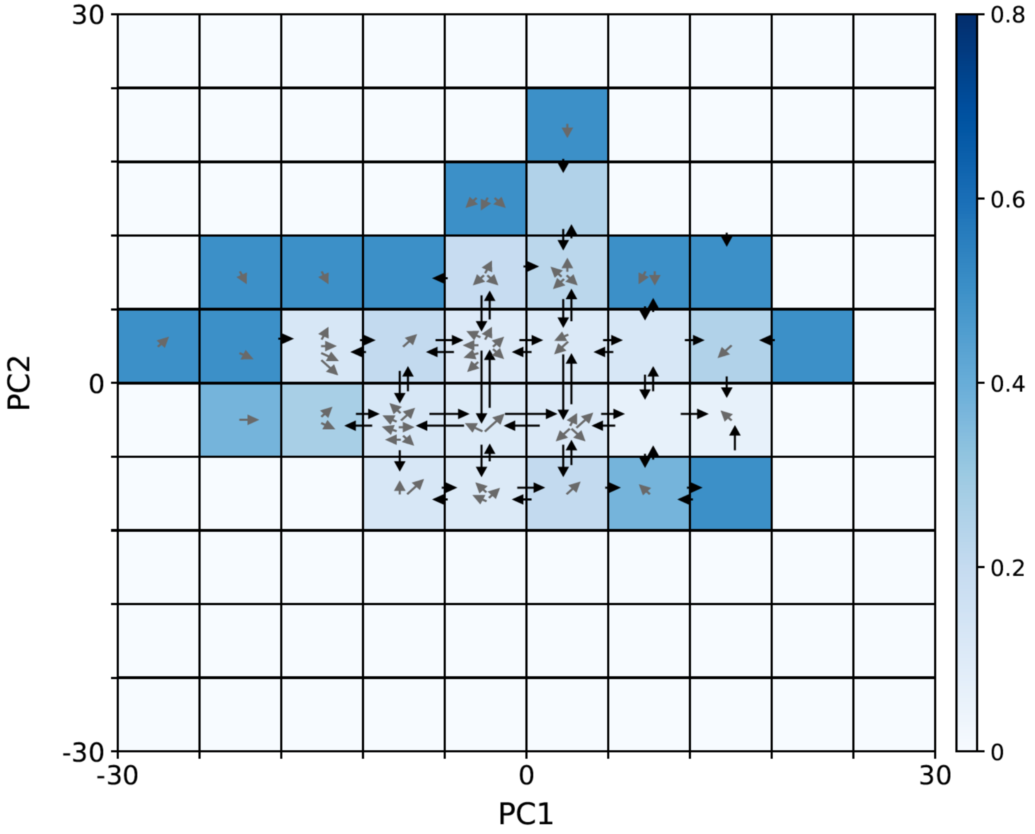


**Fig. S3. Detailed balance transition map.** Heat map of the binomial statistics-based probability of the observed ratio of forward and reverse transitions occurring by random chance. Transitions between adjacent bins are shown in black; transitions between non-adjacent bins are shown in grey.

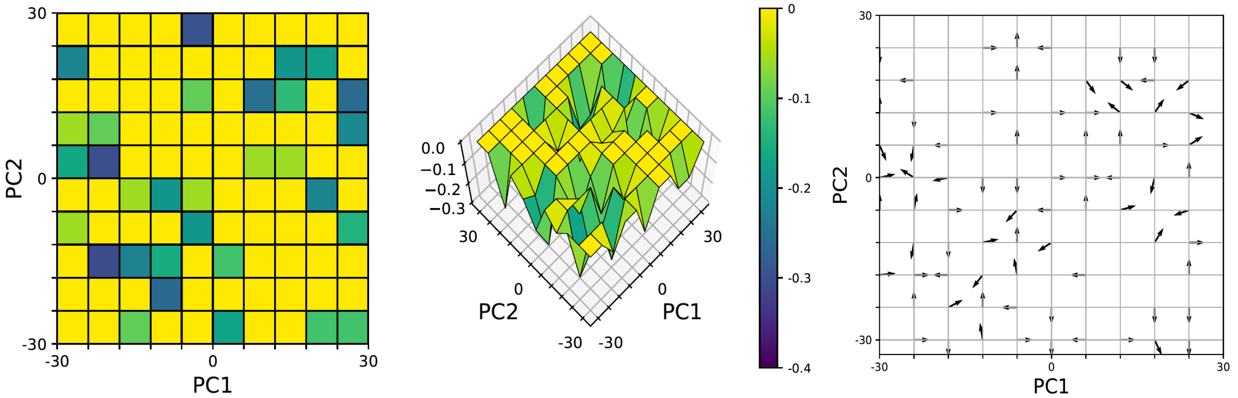


**Fig. S4. Scrambled WT MEF effective energy landscape.** Heat map (left) and surface plot (center) of a scrambled version of the effective energy landscape of WT MEF morphology space. Energy values are scaled to [-0.4, 0]. Transition vector field (right) predicted by the scrambled energy landscape.

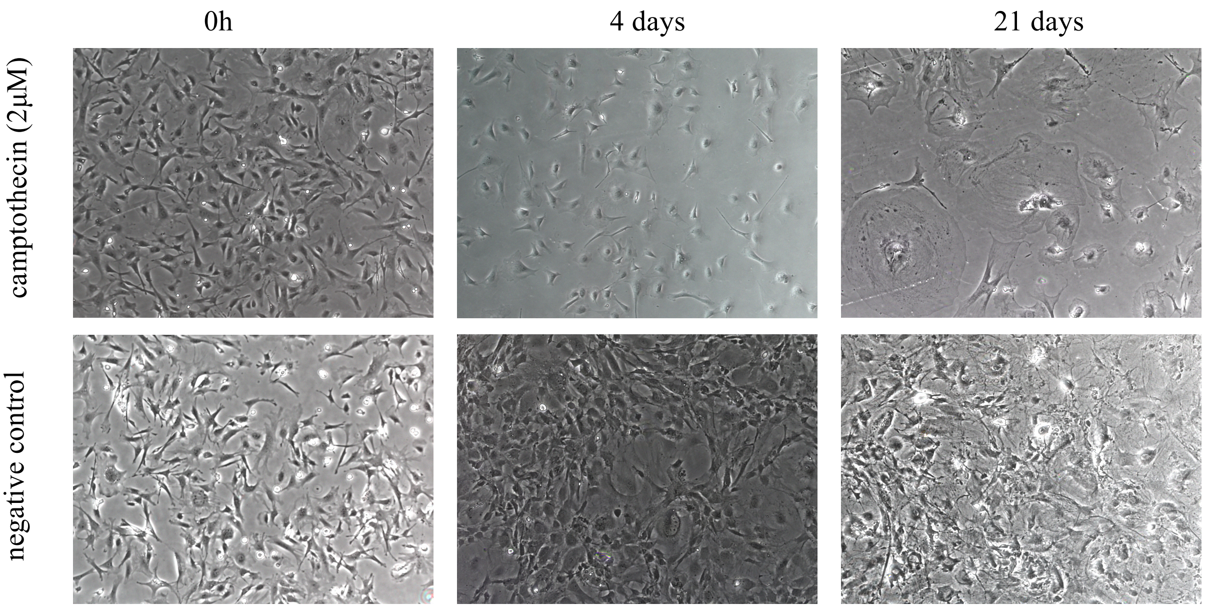


**Fig. S5. Time lapse images of WT MEFs treated with camptothecin.** Images of WT MEFs treated with 2μM camptothecin (top) and vehicle (bottom) at 0h (left), 4 days (center), and 21 days (right). Cells were incubated under either camptothecin or vehicle from 0h to 4 days; media was then replaced with drug-free media for the remaining duration of the time course.

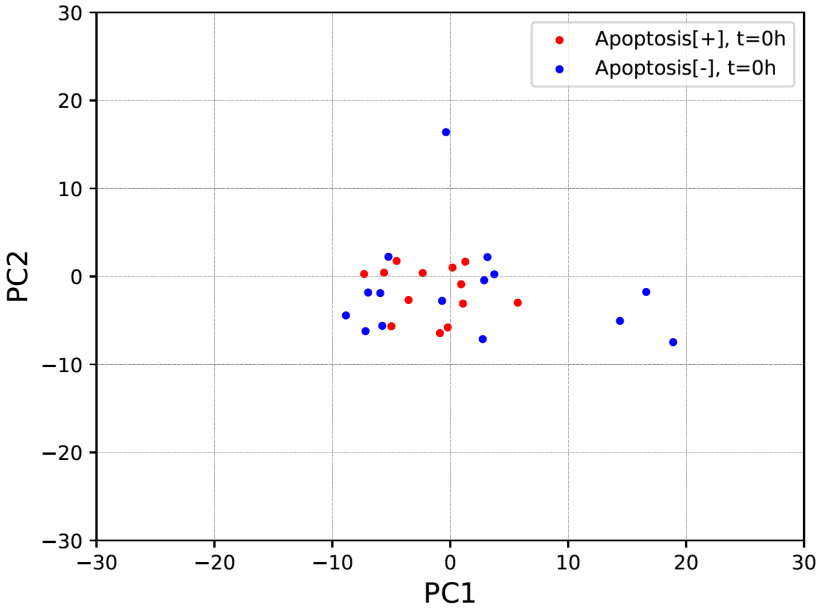


**Fig. S6. Morphology space states of Apoptosis[+] and Apoptosis[-] cells at t = 0h.** Scatter plot of single-cell coordinates of Apoptosis[+] (red) and Apoptosis[-] (blue) cells in live cell PC1 vs. PC2 space at t = 0h.

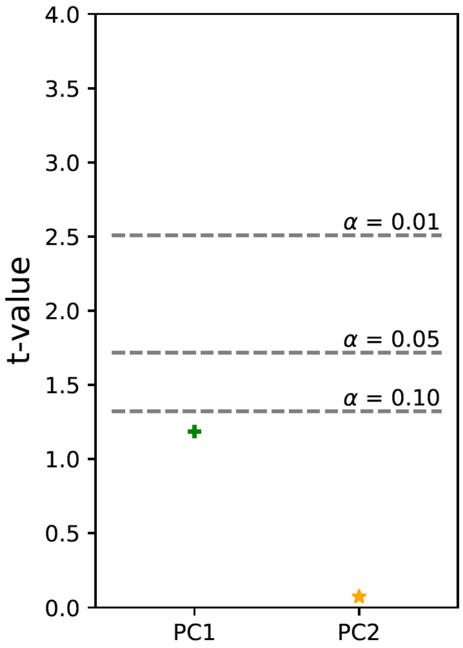


**Fig. S7. Two-sample t-test comparing PC1 and PC2 coordinates of Apoptosis[+] and Apoptosis[-] cells at t = 0h.** Two-sample t-values comparing the PC1 (green, "+") and PC2 (orange, "*") coordinates of Apoptosis[+] and Apoptosis[-] cells at t = 0h.

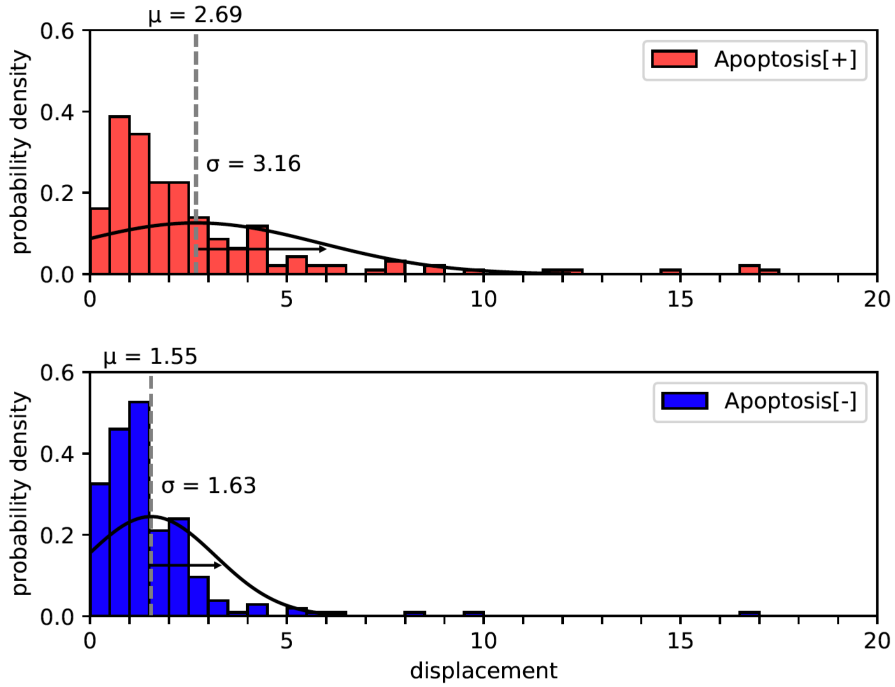


**Fig. S8: Magnitudes of transition vectors of Apoptosis[+] and Apoptosis[-] cells.** Histograms of the magnitudes of transition vectors (origin: $PC1_{(t)}$, $PC2_{(t)}$; terminus: $PC1_{(t+4h)}$, $PC2_{(t+4h)}$) of Apoptosis[+] (top, red) and Apoptosis[-] (bottom, blue) cells. Sample means (μ) are indicated by dashed grey lines. Sample standard deviations (σ) are indicated by black arrowheads. Normal distribution curves based on sample means (μ) and standard deviations (σ) are plotted in black.

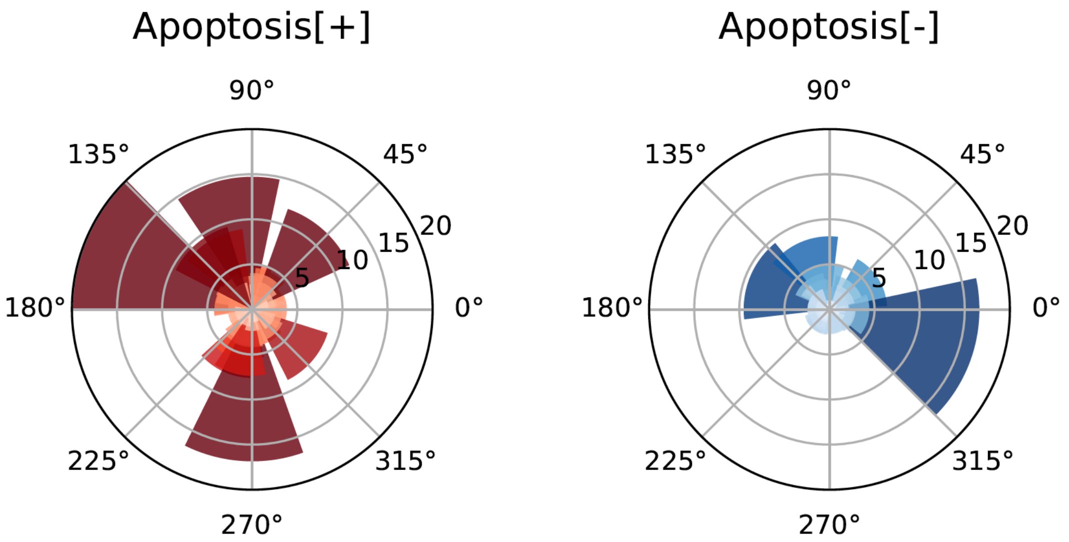


**Fig. S9: Angles of transition vectors of Apoptosis[+] and Apoptosis[-] cells.** Polar plots of the angles of transition vectors (origin: $PC1_{(t)}$, $PC2_{(t)}$; terminus: $PC1_{(t+4h)}$, $PC2_{(t+4h)}$) of Apoptosis[+] (left, red) and Apoptosis[-] (right, blue) cells. Relative to the origin (0, 0) in live cell transitions in PC1 vs. PC2 space, 0° points in the direction of (30, 0), 90° points in the direction of (0, 30), 180° points in the direction of (-30, 0), and 270° points in the direction of (0, -30). The radius of each slice correlates directly with the magnitude of the corresponding transition vector.

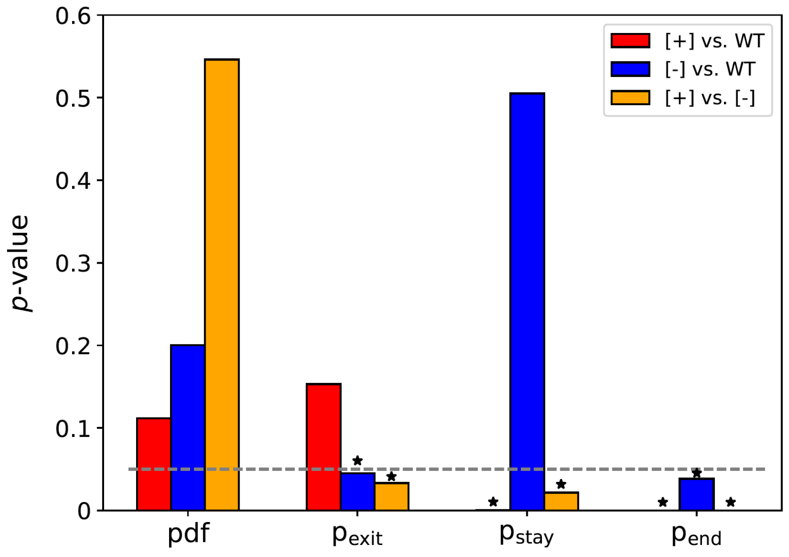


**Fig. S10: K-S plot comparing *p*-values.** Bar plot of Kolmogorov-Smirnov significance values comparing the probability distributions of WT, Apoptosis[+], and Apoptosis[-] cells. The K-S significance value at $\alpha = 0.05$ is indicated by a dashed line and statistically significant comparisons are indicated by asterisks

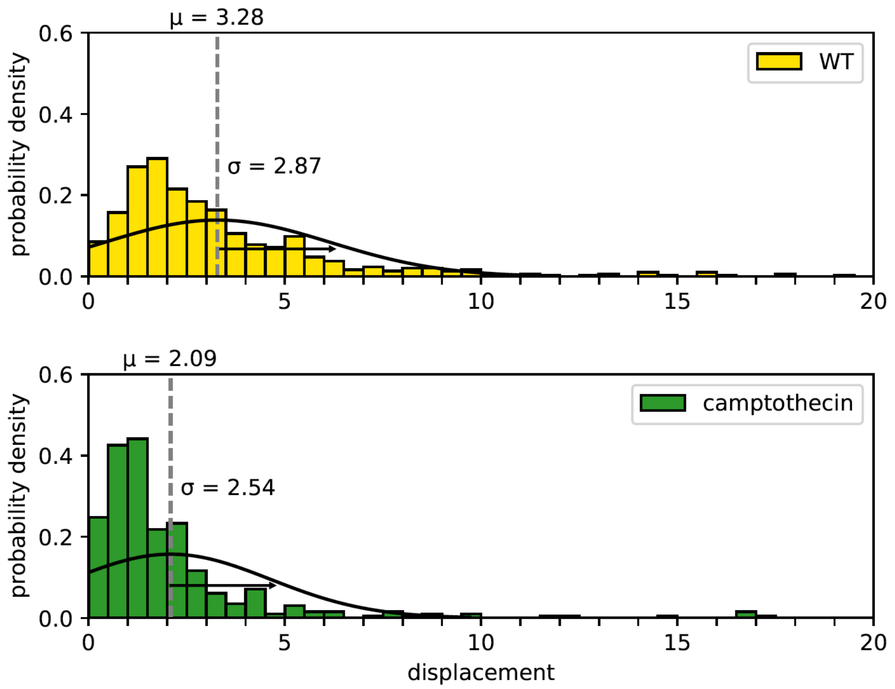


**Fig. S11: Magnitudes of transition vectors of untreated and camptothecin-treated WT MEFs.** Histograms of the magnitudes of transition vectors (origin: $PC1_{(t)}$, $PC2_{(t)}$; terminus: $PC1_{(t+4h)}$, $PC2_{(t+4h)}$) of untreated (top, yellow) and camptothecin-treated (bottom, green) WT MEFs. Sample means (μ) are indicated by dashed grey lines. Sample standard deviations (σ) are indicated by black arrowheads. Normal distribution curves based on sample means (μ) and standard deviations (σ) are plotted in black.

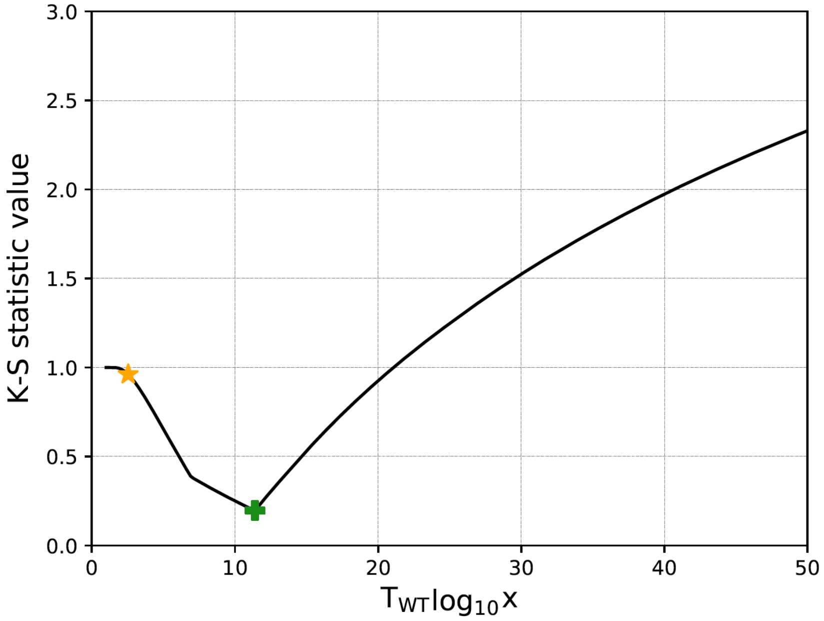


**Fig. S12. Significance test comparing observed camptothecin pdf to predicted WT pdfs.** Plot of the statistic values of two-dimensional Kolmogorov-Smirnov (K-S) tests comparing the observed pdf of camptothecin-treated WT MEFs and the predicted pdf of untreated WT MEFs occupying the untreated WT MEF effective energy landscape over a range of effective temperatures. The minimum K-S statistic value was observed at $x = 11.4$ ($1.04T_{WT}$, green +), whereas the observed effective temperature of camptothecin-treated WT MEFs was at $x = 2.54$ ($0.41T_{WT}$, orange *).

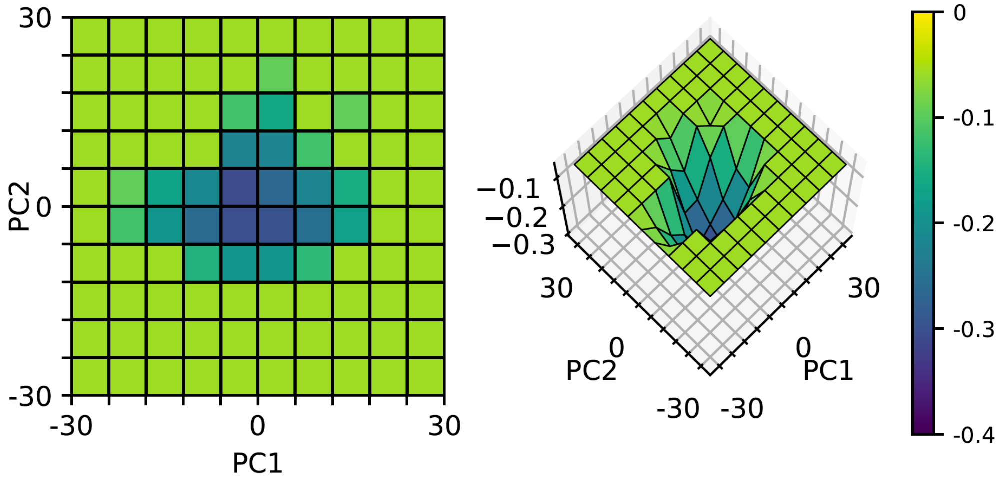


**Fig. S13. Modified WT MEF effective energy landscape.** Heat map (left) and surface plot (right) of the modified version of the effective energy landscape of WT MEF morphology space. Energy values are scaled to [-0.4, 0].

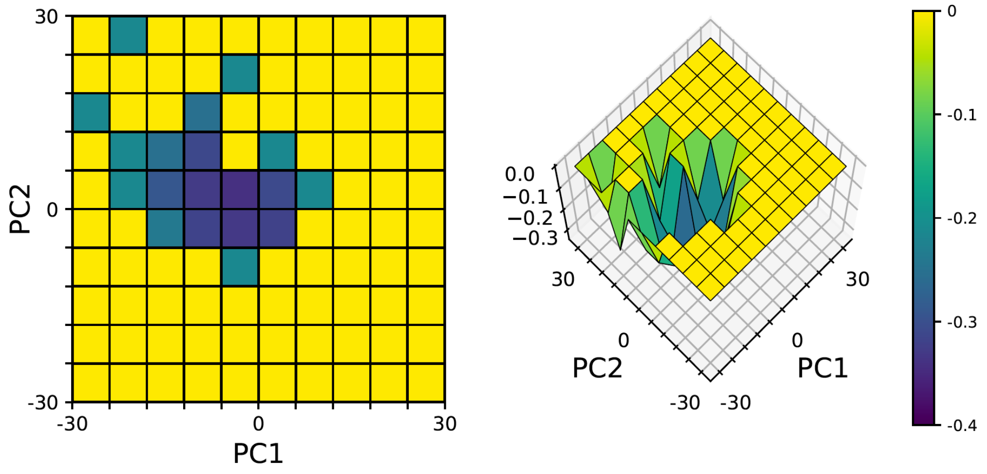


**Fig. S14. Apoptosis[+] effective energy landscape.** Heat map (left) and surface plot (right) of the effective energy landscape of Apoptosis[+] morphology space. Energy values are scaled to [-0.4, 0].

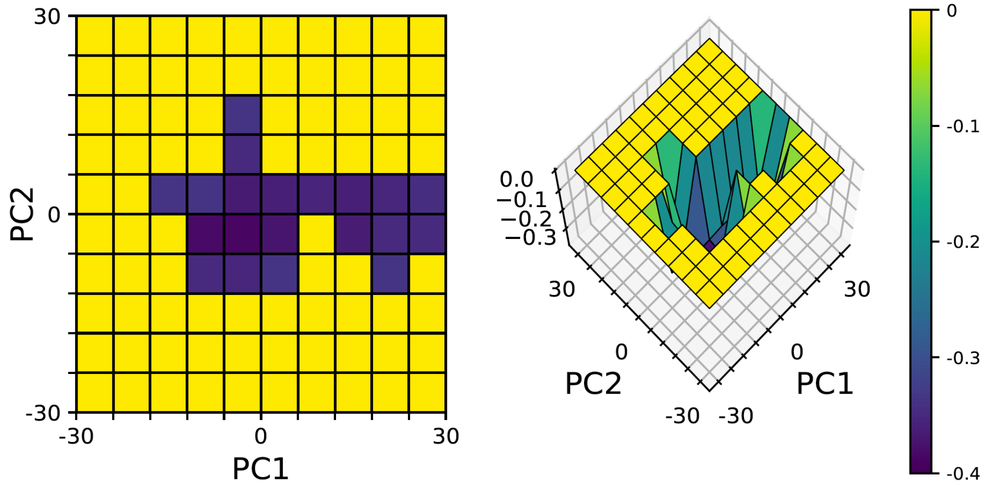


**Fig. S15. Apoptosis[-] effective energy landscape.** Heat map (left) and surface plot (right) of the effective energy landscape of Apoptosis[-] morphology space. Energy values are scaled to [-0.4, 0].

**Table S1**. **Primer sequences used for Gibson Assembly of the fluorescent reporter lentivirus expression vector.**

| **Primer Name** | **Sequence** |
|---|---|
| F-H2B_KOZAK | 5’-agcctcgaggtttaaactacgggatccggtaccgccaccatgcca-3’ |
| R-mIFP_EXTENSION | 3’-tcgaaacgtgtcagagtcaggtttccttcgcctcgatgattgaagtcg…<br>gacgacttcgtccgacctctgcacctcctcttgggacctgga-5’ |
| F-MITO | 5’-tggaggagaaccctggaccttccgtcctgacgccgc-3’ |
| R-tdTomatoLinker | 3’-gccgtggcggaggaggctcctgttgttgtaccgg-5’ |
| F-tdTomatoLinker | 5’-cggcaccgcctcctccgaggacaacaacatggcc-3’ |
| R-MITO | 3’-gcggtggtggacaaggacatgccgtacctgctcgacatgttcccttc…<br>gcctcgatgattgaagtcggacgac-5’ |
| F-EGFP_EXTENSION | 5’-ggaagcggagctactaacttcagcctgctgaagcaggctggagac…<br>gtggaggagaaccctggacctgtgagcaagggcgaggagc-3’ |

**Table S2. Apoptosis drugs screened for heterogeneous responses in WT MEFs.**

| **Drug** | | | | | | | |
|---|---|---|---|---|---|---|---|
| Actinomycin D | 200µM | 100µM | 20µM | 10µM | 4µM | 2µM | 1µM |
| Camptothecin | 40µM | 20µM | 4µM | 2µM | 800nM | 400nM | 200nM |
| Cycloheximide | 2mM | 1mM | 200µM | 100µM | 40µM | 20µM | 10µM |
| Dexamethasone | 200µM | 100µM | 20µM | 10µM | 4µM | 2µM | 1µM |
| Etoposide | 2mM | 1mM | 200µM | 100µM | 40µM | 20µM | 10µM |

**Table S3. Morphometric and Textural Feature Names and Descriptions**

| # | Feature name | Type | Description |
|---|---|---|---|
| 1 | ‘CellArea’ | cell morphometry | area of the cell (in pixels) |
| 2 | ‘CellConvexArea’ | cell morphometry | area enclosed by a convex hull around the cell |
| 3 | ‘CellEccentricity’ | cell morphometry | cell eccentricity |
| 4 | 'CellEquivDiameterArea' | cell morphometry | diameter of a circle with the same area as the cell |
| 5 | 'CellMajorAxisLength' | cell morphometry | length of the major axis of the cell |
| 6 | 'CellMinorAxisLength' | cell morphometry | length of the minor axis of the cell |
| 7 | 'CellPerimeter’ | cell morphometry | perimeter of the cell |
| 8 | 'CellSolidity' | cell morphometry | proportion of pixels contained by the cell convex hull that are also in the cell |
| 9 | 'CellConvexPerimeter' | cell morphometry | perimeter of a convex hull around the cell |
| 10 | 'CellEquivDiameterPerim' | cell morphometry | diameter of a circle with the same perimeter length as the cell |
| 11 | 'CellRatioMajorMinor' | cell morphometry | (cell major axis length) / (cell minor axis length) |
| 12 | 'CellCircularity' | cell morphometry | (cell area) / (area of a circle with diameter equal to the major axis length of the cell) |
| 13 | 'CellMeanCentroidDist' | cell morphometry | mean distance between the cell centroid and a point on its perimeter |
| 14 | 'CellMaxCentroidDist' | cell morphometry | maximum distance between the cell centroid and a point on its perimeter |
| 15 | 'CellMinCentroidDist' | cell morphometry | maximum distance between the cell centroid and a point on its perimeter |
| 16 | 'CellCVCentroidDist' | cell morphometry | coefficient of variation of distances between the cell centroid and all points on its perimeter |
| 17 | 'CellVarCentroidDist' | cell morphometry | variance of distances between the cell centroid and all points on its perimeter |
| 18 | 'CellLowSmoothArea' | cell morphometry | area of cell after fine-grain smoothing |
| 19 | 'CellLowSmoothPerimeter' | cell morphometry | perimeter of cell after fine-grain smoothing |
| 20 | 'CellLowSmoothOrigArea Change' | cell morphometry | (cell area after fine-grain smoothing) – (cell area) |
| 21 | 'CellLowSmoothOrigArea ChangeRel' | cell morphometry | [(cell area after fine-grain smoothing) – (cell area)] / (cell area) |
| 22 | 'CellLowSmoothOrig PerimeterChange' | cell morphometry | (cell perimeter after fine-grain smoothing) – (cell perimeter) |
| 23 | 'CellLowSmoothOrig PerimeterChangeRel' | cell morphometry | [(cell perimeter after fine-grain smoothing) – (cell perimeter)] / (cell perimeter) |
| 24 | 'CellLowSmoothOrigArea Ratio' | cell morphometry | (cell area after fine-grain smoothing) / (cell area) |
| 25 | 'CellLowSmoothOrig PerimeterRatio' | cell morphometry | (cell perimeter after fine-grain smoothing) / (cell perimeter) |
| 26 | 'CellHighSmoothArea' | cell morphometry | area of cell after coarse-grain smoothing |
| 27 | 'CellHighSmoothPerimeter ' | cell morphometry | perimeter of cell after coarse-grain smoothing |
| 28 | 'CellHighSmoothOrigArea | cell morphometry | (cell area after coarse-grain smoothing) - |

| | | | |
|---|---|---|---|
| | Change' | | (cell area) |
| 29 | 'CellHighSmoothOrigArea ChangeRel' | cell morphometry | [(cell area after coarse-grain smoothing) – (cell area)] / (cell area) |
| 30 | 'CellHighSmoothOrig PerimeterChange' | cell morphometry | (cell perimeter after coarse-grain smoothing) – (cell perimeter) |
| 31 | 'CellHighSmoothOrig PerimeterChangeRel' | cell morphometry | [(cell perimeter after coarse-grain smoothing) – (cell perimeter)] / (cell perimeter) |
| 32 | 'CellHighSmoothOrigArea Ratio' | cell morphometry | (cell area after coarse-grain smoothing) / (cell area) |
| 33 | 'CellHighSmoothOrig PerimeterRatio' | cell morphometry | (cell perimeter after coarse-grain smoothing) / (cell perimeter) |
| 34 | 'NucNumber' | nuc morphometry | number of nuclei |
| 35 | 'NucArea' | nuc morphometry | total area of nuclei |
| 36 | 'NucMeanArea' | nuc morphometry | (total area of nuclei) / (number of nuclei) |
| 37 | 'NucCellAreaRatio' | nuc morphometry | (area of nuclei) / (cell area) |
| 38 | 'NucCellCentroidDist' | nuc morphometry | distance between nucleus and cell centroids |
| 39 | 'NucConvexArea' | nuc morphometry | area enclosed by a convex hull around the nucleus |
| 40 | 'NucEccentricity' | nuc morphometry | nucleus eccentricity |
| 41 | 'NucEquivDiameterArea' | nuc morphometry | diameter of a circle with the same area as the nucleus |
| 42 | 'NucMajorAxisLength' | nuc morphometry | length of the major axis of the nucleus |
| 43 | 'NucMinorAxisLength' | nuc morphometry | length of the minor axis of the nucleus |
| 44 | 'NucPerimeter' | nuc morphometry | perimeter of the nucleus |
| 45 | 'NucMeanPerimeter' | nuc morphometry | (perimeter of the nucleus) / (number of nuclei) |
| 46 | 'NucCellPerimeterRatio' | nuc morphometry | (nucleus perimeter) / (cell perimeter) |
| 47 | 'NucSolidity' | nuc morphometry | proportion of pixels contained by the nucleus convex hull that are also in the nucleus |
| 48 | 'NucConvexPerimeter' | nuc morphometry | perimeter of a convex hull around the nucleus |
| 49 | 'NucEquivDiameterPerim' | nuc morphometry | diameter of a circle with the same perimeter length as the nucleus |
| 50 | 'NucRatioMajorMinor' | nuc morphometry | (nucleus major axis length) / (nucleus minor axis length) |
| 51 | 'NucCircularity' | nuc morphometry | (nucleus area) / (area of a circle with diameter equal to the major axis length of the nucleus) |
| 52 | 'NucMeanCentroidDist' | nuc morphometry | mean distance between the nucleus centroid and a point on its perimeter |
| 53 | 'NucMaxCentroidDist' | nuc morphometry | maximum distance between the nucleus centroid and a point on its perimeter |
| 54 | 'NucMinCentroidDist' | nuc morphometry | minimum distance between the nucleus centroid and a point on its perimeter |
| 55 | 'NucCVCentroidDist' | nuc morphometry | coefficient of variation of distances between the nucleus centroid and all points on its perimeter |
| 56 | 'NucVarCentroidDist' | nuc morphometry | variance of distances between the nucleus centroid and all points on its perimeter |

| | | | |
|---|---|---|---|
| 57 | 'NucLowSmoothArea' | nuc morphometry | area of nucleus after fine-grain smoothing |
| 58 | 'NucLowSmoothPerimeter' | nuc morphometry | perimeter of nucleus after fine-grain smoothing |
| 59 | 'NucLowSmoothOrigArea Change' | nuc morphometry | (nucleus area after fine-grain smoothing) – (nucleus area) |
| 60 | 'NucLowSmoothOrigArea ChangeRel' | nuc morphometry | [(nucleus area after fine-grain smoothing) – (nucleus area)] / (nucleus area) |
| 61 | 'NucLowSmoothOrig PerimeterChange' | nuc morphometry | (nucleus perimeter after fine-grain smoothing) – (nucleus perimeter) |
| 62 | 'NucLowSmoothOrig PerimeterChangeRel' | nuc morphometry | [(nucleus perimeter after fine-grain smoothing) – (nucleus perimeter)] / (nucleus perimeter) |
| 63 | 'NucLowSmoothOrigArea Ratio' | nuc morphometry | (nucleus area after fine-grain smoothing) / (nucleus area) |
| 64 | 'NucLowSmoothOrig PerimeterRatio' | nuc morphometry | (nucleus perimeter after fine-grain smoothing) / (nucleus perimeter) |
| 65 | 'NucHighSmoothArea' | nuc morphometry | area of nucleus after coarse-grain smoothing |
| 66 | 'NucHighSmoothPerimeter ' | nuc morphometry | perimeter of nucleus after coarse-grain smoothing |
| 67 | 'NucHighSmoothOrigArea Change' | nuc morphometry | (nucleus area after coarse-grain smoothing) - (nucleus area) |
| 68 | 'NucHighSmoothOrigArea ChangeRel' | nuc morphometry | [(nucleus area after coarse-grain smoothing) – (nucleus area)] / (nucleus area) |
| 69 | 'NucHighSmoothOrig PerimeterChange' | nuc morphometry | (nucleus perimeter after coarse-grain smoothing) – (nucleus perimeter) |
| 70 | 'NucHighSmoothOrig PerimeterChangeRel' | nuc morphometry | [(nucleus perimeter after coarse-grain smoothing) – (nucleus perimeter)] / (nucleus perimeter) |
| 71 | 'NucHighSmoothOrig AreaRatio' | nuc morphometry | (nucleus area after coarse-grain smoothing) / (nucleus area) |
| 72 | NucHighSmoothOrig PerimeterRatio' | nuc morphometry | (nucleus perimeter after coarse-grain smoothing) / (nucleus perimeter) |
| 73 | 'CellMeanContrast' | cell texture | mean cell contrast (a measure of the intensity contrast between a pixel and its neighbor); contrast is 0 for a constant image |
| 74 | 'CellMinContrast' | cell texture | minimum cell contrast |
| 75 | 'CellMaxContrast' | cell texture | maximum cell contrast |
| 76 | 'CellCVContrast' | cell texture | coefficient of variation of cell contrast values |
| 77 | 'CellVarContrast' | cell texture | variance of cell contrast values |
| 78 | 'CellMeanCorrelation' | cell texture | mean cell correlation (a measure of how correlated a pixel is to its neighbor) |
| 79 | 'CellMinCorrelation' | cell texture | minimum cell correlation |
| 80 | 'CellMaxCorrelation' | cell texture | maximum cell correlation |
| 81 | 'CellCVCorrelation' | cell texture | coefficient of variation of cell correlation values |
| 82 | 'CellVarCorrelation' | cell texture | variance of cell correlation values |
| 83 | 'CellMeanEnergy' | cell texture | mean cell energy (the sum of squared elements in the mitochondria gray-level co-occurrence matrix); energy is 1 for a constant image |

| | | | |
|---|---|---|---|
| 84 | 'CellMinEnergy' | cell texture | minimum cell energy |
| 85 | 'CellMaxEnergy' | cell texture | maximum cell energy |
| 86 | 'CellCVEnergy' | cell texture | coefficient of variation of cell energy values |
| 87 | 'CelVarEnergy' | cell texture | variance of cell energy values |
| 88 | 'CellMeanHomogeneity' | cell texture | mean cell homogeneity (a measure of how close the distribution of values in the gray-level co-occurrence matrix is to the diagonal of the matrix); homogeneity is 1 for a diagonal grey-level co-occurrence matrix |
| 89 | 'CellMinHomogeneity' | cell texture | minimum cell homogeneity |
| 90 | 'CellMaxHomogeneity' | cell texture | maximum cell homogeneity |
| 91 | 'CellCVHomogeneity' | cell texture | coefficient of variation of cell homogeneity values |
| 92 | 'CellVarHomogeneity' | cell texture | variance of cell homogeneity values |
| 93 | 'CellFracTotalDisc1' | cell texture | fraction of nucleus intensity located within ¼ $radius_{cell}$ of the nucleus centroid |
| 94 | 'CellCVDisc1' | cell texture | coefficient of variation of intensity values of cell located within ¼ $radius_{cell}$ of the nucleus centroid |
| 95 | 'CellVarDisc1' | cell texture | variance of intensity values of cell located within ¼ $radius_{cell}$ of the nucleus centroid |
| 96 | 'CellFracTotalDisc2' | cell texture | fraction of nucleus intensity located within ½ $radius_{cell}$ of the nucleus centroid |
| 97 | 'CellCVDisc2' | cell texture | coefficient of variation of intensity values of cell located within ½ $radius_{cell}$ of the nucleus centroid |
| 98 | 'CellVarDisc2' | cell texture | variance of intensity values of cell located within ½ $radius_{cell}$ of the nucleus centroid |
| 99 | 'CellFracTotalDisc3' | cell texture | fraction of nucleus intensity located within ¾ $radius_{cell}$ of the nucleus centroid |
| 100 | 'CellCVDisc3' | cell texture | coefficient of variation of intensity values of cell located within ¾ $radius_{cell}$ of the nucleus centroid |
| 101 | 'CellVarDisc3' | cell texture | variance of intensity values of cell located within ¾ $radius_{cell}$ of the nucleus centroid |
| 102 | 'CellFracTotalDisc4' | cell texture | fraction of nucleus intensity located within $radius_{cell}$ of the nucleus centroid |
| 103 | 'CellCVDisc4' | cell texture | coefficient of variation of intensity values of cell located within $radius_{cell}$ of the nucleus centroid |
| 104 | 'CellVarDisc4' | cell texture | variance of intensity values of cell located within $radius_{cell}$ of the nucleus centroid |
| 105 | 'NucMeanContrast' | nuc texture | mean nucleus contrast (a measure of the intensity contrast between a pixel and its neighbor); contrast is 0 for a constant image |
| 106 | 'NucMinContrast' | nuc texture | minimum nucleus contrast |
| 107 | 'NucMaxContrast' | nuc texture | maximum nucleus contrast |
| 108 | 'NucCVContrast' | nuc texture | coefficient of variation of nucleus contrast values |
| 109 | 'NucVarContrast' | nuc texture | variance of nucleus contrast values |

| | | | |
|---|---|---|---|
| 110 | 'NucMeanCorrelation' | nuc texture | mean nucleus correlation (a measure of how correlated a pixel is to its neighbor) |
| 111 | 'NucMinCorrelation' | nuc texture | minimum nucleus correlation |
| 112 | 'NucMaxCorrelation' | nuc texture | maximum nucleus correlation |
| 113 | 'NucCVCorrelation' | nuc texture | coefficient of variation of nucleus correlation values |
| 114 | 'NucVarCorrelation' | nuc texture | variance of nucleus correlation values |
| 115 | 'NucMeanEnergy' | nuc texture | mean nucleus energy (the sum of squared elements in the mitochondria gray-level co-occurrence matrix); energy is 1 for a constant image |
| 116 | 'NucMinEnergy' | nuc texture | minimum nucleus energy |
| 117 | 'NucMaxEnergy' | nuc texture | maximum nucleus energy |
| 118 | 'NucCVEnergy' | nuc texture | coefficient of variation of nucleus energy values |
| 119 | 'NucVarEnergy' | nuc texture | variance of nucleus energy values |
| 120 | 'NucMeanHomogeneity' | nuc texture | mean nucleus homogeneity (a measure of how close the distribution of values in the gray-level co-occurrence matrix is to the diagonal of the matrix); homogeneity is 1 for a diagonal grey-level co-occurrence matrix |
| 121 | 'NucMinHomogeneity' | nuc texture | minimum nucleus homogeneity |
| 122 | 'NucMaxHomogeneity' | nuc texture | maximum nucleus homogeneity |
| 123 | 'NucCVHomogeneity' | nuc texture | coefficient of variation of nucleus homogeneity values |
| 124 | 'NucVarHomogeneity' | nuc texture | variance of nucleus homogeneity values |
| 125 | 'NucFracTotalDisc1' | nuc texture | fraction of nucleus intensity located within ¼ $radius_{nucleus}$ of the nucleus centroid |
| 126 | 'NucCVDisc1' | nuc texture | coefficient of variation of intensity values of nucleus located within ¼ $radius_{nucleus}$ of the nucleus centroid |
| 127 | 'NucVarDisc1' | nuc texture | variance of intensity values of nucleus located within ¼ $radius_{nucleus}$ of the nucleus centroid |
| 128 | 'NucFracTotalDisc2' | nuc texture | fraction of nucleus intensity located within ½ $radius_{nucleus}$ of the nucleus centroid |
| 129 | 'NucCVDisc2' | nuc texture | coefficient of variation of intensity values of nucleus located within ½ $radius_{nucleus}$ of the nucleus centroid |
| 130 | 'NucVarDisc2' | nuc texture | variance of intensity values of nucleus located within ½ $radius_{nucleus}$ of the nucleus centroid |
| 131 | 'NucFracTotalDisc3' | nuc texture | fraction of nucleus intensity located within ¾ $radius_{nucleus}$ of the nucleus centroid |
| 132 | 'NucCVDisc3' | nuc texture | coefficient of variation of intensity values of nucleus located within ¾ $radius_{nucleus}$ of the nucleus centroid |
| 133 | 'NucVarDisc3' | nuc texture | variance of intensity values of nucleus located within ¾ $radius_{nucleus}$ of the nucleus centroid |

| | | | |
|---|---|---|---|
| 134 | 'NucFracTotalDisc4' | nuc texture | fraction of nucleus intensity located within $radius_{nucleus}$ of the nucleus centroid |
| 135 | 'NucCVDisc4' | nuc texture | coefficient of variation of intensity values of nucleus located within $radius_{nucleus}$ of the nucleus centroid |
| 136 | 'NucVarDisc4' | nuc texture | variance of intensity values of nucleus located within $radius_{nucleus}$ of the nucleus centroid |
| 137 | 'MitoNumber' | mito morphometry | number of mitochondria |
| 138 | 'MitoSumArea' | mito morphometry | total area of mitochondria objects |
| 139 | 'MitoMeanArea' | mito morphometry | mean mitochondria area |
| 140 | 'MitoMedianArea' | mito morphometry | median mitochondria area |
| 141 | 'MitoMaxArea' | mito morphometry | maximum mitochondria area |
| 142 | 'MitoMinArea' | mito morphometry | minimum mitochondria area |
| 143 | 'MitoCVArea' | mito morphometry | coefficient of variation of all mitochondria area measurements |
| 144 | 'MitoVarArea' | mito morphometry | variance of all mitochondria area measurements |
| 145 | 'MitoCellAreaRatio' | mito morphometry | (total area of mitochondria) / (cell area) |
| 146 | 'MitoSumPerimeter' | mito morphometry | total perimeter of mitochondria objects |
| 147 | 'MitoMeanPerimeter' | mito morphometry | mean mitochondria perimeter |
| 148 | 'MitoMedianPerimeter' | mito morphometry | median mitochondria perimeter |
| 149 | 'MitoMaxPerimeter' | mito morphometry | maximum mitochondria perimeter |
| 150 | 'MitoMinPerimeter' | mito morphometry | minimum mitochondria perimeter |
| 151 | 'MitoCVPerimeter' | mito morphometry | coefficient of variation of all mitochondria perimeter measurements |
| 152 | 'MitoVarPerimeter' | mito morphometry | variance of all mitochondria perimeter measurements |
| 153 | 'MitoCellPerimeterRatio' | mito morphometry | (total perimeter of mitochondria) / (cell perimeter) |
| 154 | 'MitoCellCentroidDist' | mito morphometry | distance between mitochondria and cell centroids |
| 155 | 'MitoNucCentroidDist' | mito morphometry | distance between mitochondria and nucleus centroids |
| 156 | 'MitoObjSkelRatio' | mito morphometry | (total area of mitochondria) / (total area of mitochondria skeleton); represents the average width of mitochondria |
| 157 | 'MitoNumBranchpoints' | mito | number of mitochondria branchpoints |

| | | | |
|---|---|---|---|
| | | morphometry | |
| 158 | 'MitoNumEndpoints' | mito morphometry | number of mitochondria endpoints |
| 159 | 'MitoNumberFused' | mito morphometry | number of fused mitochondria (contain a branchpoint) |
| 160 | 'MitoMeanFusedArea' | mito morphometry | mean area of fused mitochondria |
| 161 | 'MitoMaxFusedArea' | mito morphometry | maximum area of fused mitochondria |
| 162 | 'MitoMinFusedArea' | mito morphometry | minimum area of fused mitochondria |
| 163 | 'MitoCVFusedArea' | mito morphometry | coefficient of variation of all fused mitochondria area measurements |
| 164 | 'MitoVarFusedArea' | mito morphometry | variance of all fused mitochondria area measurements |
| 165 | 'MitoPercentFusedTotalArea' | mito morphometry | (area of fused mitochondria) / (total area of all mitochondria) |
| 166 | 'MitoNumberFragmented' | mito morphometry | number of fragmented mitochondria (does not contain a branchpoint) |
| 167 | 'MitoMeanFragmentedArea' | mito morphometry | mean area of fragmented mitochondria |
| 168 | 'MitoMaxFragmentedArea' | mito morphometry | maximum area of fragmented mitochondria |
| 169 | 'MitoMinFragmentedArea' | mito morphometry | minimum area of fragmented mitochondria |
| 170 | 'MitoCVFragmentedArea' | mito morphometry | coefficient of variation of all fragmented mitochondria area measurements |
| 171 | 'MitoVarFragmentedArea' | mito morphometry | variance of variation of all fragmented mitochondria area measurements |
| 172 | 'MitoRatioFusedFragmented' | mito morphometry | (area of fused mitochondria) / (area of fragmented mitochondria) |
| 173 | 'MitoPercentFragmented TotalArea' | mito morphometry | (area of fragmented mitochondria) / (total area of all mitochondria) |
| 174 | 'MitoMeanContrast' | mito texture | mean mitochondria contrast (a measure of the intensity contrast between a pixel and its neighbor); contrast is 0 for a constant image |
| 175 | 'MitoMinContrast' | mito texture | minimum mitochondria contrast |
| 176 | ‘MitoMaxContrast' | mito texture | maximum mitochondria contrast |
| 177 | 'MitoCVContrast' | mito texture | coefficient of variation of mitochondria contrast values |
| 178 | 'MitoVarContrast' | mito texture | variance of mitochondria contrast values |
| 179 | 'MitoMeanCorrelation' | mito texture | mean mitochondria correlation (a measure of how correlated a pixel is to its neighbor) |
| 180 | 'MitoMinCorrelation' | mito texture | minimum mitochondria correlation |
| 181 | 'MitoMaxCorrelation' | mito texture | maximum mitochondria correlation |
| 182 | 'MitoCVCorrelation' | mito texture | coefficient of variation of mitochondria correlation values |
| 183 | 'MitoVarCorrelation' | mito texture | variance of mitochondria correlation values |
| 184 | 'MitoMeanEnergy' | mito texture | mean mitochondria energy (the sum of squared elements in the mitochondria gray- |

| | | | |
|---|---|---|---|
| | | | level co-occurrence matrix); energy is 1 for a constant image |
| 185 | 'MitoMinEnergy' | mito texture | minimum mitochondria energy |
| 186 | 'MitoMaxEnergy' | mito texture | maximum mitochondria energy |
| 187 | 'MitoCVEnergy' | mito texture | coefficient of variation of mitochondria energy values |
| 188 | 'MitoVarEnergy' | mito texture | variance of mitochondria energy values |
| 189 | 'MitoMeanHomogeneity' | mito texture | mean mitochondria homogeneity (a measure of how close the distribution of values in the gray-level co-occurrence matrix is to the diagonal of the matrix); homogeneity is 1 for a diagonal grey-level co-occurrence matrix |
| 190 | 'MitoMinHomogeneity' | mito texture | minimum mitochondria homogeneity |
| 191 | 'MitoMaxHomogeneity' | mito texture | maximum mitochondria homogeneity |
| 192 | 'MitoCVHomogeneity' | mito texture | coefficient of variation of mitochondria homogeneity values |
| 193 | 'MitoVarHomogeneity' | mito texture | variance of mitochondria homogeneity values |
| 194 | 'MitoFracTotalDisc1' | mito texture | fraction of mitochondria intensity located within ¼ $radius_{mitochondria}$ of the nucleus centroid |
| 195 | 'MitoCVDisc1' | mito texture | coefficient of variation of intensity values of mitochondria located within ¼ $radius_{mitochondria}$ of the nucleus centroid |
| 196 | 'MitoVarDisc1' | mito texture | variance of intensity values of mitochondria located within ¼ $radius_{mitochondria}$ of the nucleus centroid |
| 197 | 'MitoFracTotalDisc2' | mito texture | fraction of mitochondria intensity located within ½ $radius_{mitochondria}$ of the nucleus centroid |
| 198 | 'MitoCVDisc2' | mito texture | coefficient of variation of intensity values of mitochondria located within ½ $radius_{mitochondria}$ of the nucleus centroid |
| 199 | 'MitoVarDisc2' | mito texture | variance of intensity values of mitochondria located within ½ $radius_{mitochondria}$ of the nucleus centroid |
| 200 | 'MitoFracTotalDisc3' | mito texture | fraction of mitochondria intensity located within ¾ $radius_{mitochondria}$ of the nucleus centroid |
| 201 | 'MitoCVDisc3' | mito texture | coefficient of variation of intensity values of mitochondria located within ¾ $radius_{mitochondria}$ of the nucleus centroid |
| 202 | 'MitoVarDisc3' | mito texture | variance of intensity values of mitochondria located within ¾ $radius_{mitochondria}$ of the nucleus centroid |
| 203 | 'MitoFracTotalDisc4' | mito texture | fraction of mitochondria intensity located within $radius_{mitochondria}$ of the nucleus centroid |
| 204 | 'MitoCVDisc4' | mito texture | coefficient of variation of intensity values of mitochondria located within $radius_{mitochondria}$ of the nucleus centroid |

| | | | |
|---|---|---|---|
| 205 | 'MitoVarDisc4' | mito texture | variance of intensity values of mitochondria located within $radius_{mitochondria}$ of the nucleus centroid |

**Table S4**. **Dominant loadings of the first three Principal Components of fixed WT MEFs.** The top 10 loadings of the first three principal components (PCs) of a 205-feature set analysis of fixed WT MEFs. PC1 captures 29.6% of total variance and is dominated by cell and nucleus morphometric features. PC2 captures 10.6% of total variance and is dominated by cell morphometric and textural features. PC3 captures 6.7% of total variance and is dominated by mitochondrial textural and cell morphometric features.

**Principal Component 1; variance explained = 29.6%**

| Feature Name | Feature Type | Loading |
|---|---|---|
| CellEquivDiameterArea | Cell morphometry | 0.1185 |
| NucEquivDiameterArea | Nucleus morphometry | 0.1173 |
| NucMeanCentroidDist | Nucleus morphometry | 0.1166 |
| CellMeanCentroidDist | Cell morphometry | 0.1151 |
| NucMinorAxisLength | Nucleus morphometry | 0.1145 |
| NucConvexPerimeter | Nucleus morphometry | 0.1143 |
| NucHighSmoothPerimeter | Nucleus morphometry | 0.1140 |
| CellConvexPerimeter | Cell morphometry | 0.1133 |
| CellHighSmoothPerimeter | Cell morphometry | 0.1132 |
| NucLowSmoothPerimeter | Nucleus morphometry | 0.1130 |

**Principal Component 2; variance explained = 10.6%**

| Feature Name | Feature Type | Loading |
|---|---|---|
| CellCVDisc3 | Cell texture | -0.1558 |
| CellCVDisc4 | Cell texture | -0.1528 |
| CellCVDisc2 | Cell texture | -0.1526 |
| CellCVDisc1 | Cell texture | -0.1397 |
| CellFracTotalDisc1 | Cell texture | -0.1393 |
| CellCVCentroidDist | Cell texture | -0.1318 |
| CellHighSmoothOrigPerimeterChangeRel | Cell morphometry | 0.1223 |
| CellHighSmoothOrigPerimeterRatio | Cell morphometry | 0.1223 |
| CellCircularity | Cell morphometry | 0.1216 |
| CellSolidity | Cell morphometry | 0.1191 |

**Principal Component 3; variance explained = 6.7%**

| Feature Name | Feature Type | Loading |
|---|---|---|
| MitoCVDisc2 | Mitochondria texture | 0.1725 |
| MitoCVDisc3 | Mitochondria texture | 0.1716 |
| MitoCVDisc4 | Mitochondria texture | 0.1698 |
| CellCVCorrelation | Cell texture | 0.1573 |
| CellCVCentroidDist | Cell morphometry | 0.1546 |
| CellCircularity | Cell morphometry | -0.1464 |
| MitoVarCorrelation | Mitochondria texture | 0.1436 |
| CellVarCorrelation | Cell texture | 0.1433 |
| MitoCVCorrelation | Mitochondria texture | 0.1358 |
| CellVarHomogeneity | Cell texture | 0.1344 |

**Table S5**. **Dominant loadings of the first three Principal Components of live WT MEFs.** The top 10 loadings of the first three principal components (PCs) of a 205-feature set analysis of live WT MEFs. PC1 captures 23.4% of total variance and is dominated by cell and nucleus morphometric features. PC2 captures 11.8% of total variance and is dominated by cell and nucleus textural features. PC3 captures 8.6% of total variance and is dominated by nucleus morphometric and textural features.

**Principal Component 1; variance explained = 23.4%**

| Feature Name | Feature Type | Loading |
|---|---|---|
| CellMeanCentroidDist | Cell morphometry | 0.1324 |
| CellConvexPerimeter | Cell morphometry | 0.1321 |
| NucEquivDiameterArea | Nucleus morphometry | 0.1307 |
| CellEquivDiameterArea | Cell morphometry | 0.1305 |
| CellLowSmoothPerimeter | Cell morphometry | 0.1296 |
| NucMeanCentroidDist | Nucleus morphometry | 0.1295 |
| CellPerimeter | Cell morphometry | 0.1292 |
| CellEquivDiameterPerim | Cell morphometry | 0.1292 |
| CellHighSmoothPerimeter | Cell morphometry | 0.1290 |

**Principal Component 2; variance explained = 11.8%**

| Feature Name | Feature Type | Loading |
|---|---|---|
| NucMinContrast | Nucleus texture | 0.1583 |
| CellMinContrast | Cell texture | 0.1563 |
| CellMeanContrast | Cell texture | 0.1551 |
| CellMeanHomogeneity | Cell texture | -0.1538 |
| CellMaxHomogeneity | Cell texture | -0.1530 |
| CellMeanCorrelation | Cell texture | -0.1529 |
| CellMaxCorrelation | Cell texture | -0.1526 |
| CellMinHomogeneity | Cell texture | -0.1524 |
| CellMinCorrelation | Cell texture | -0.1519 |
| NucMaxHomogeneity | Nucleus texture | -0.1515 |

**Principal Component 3; variance explained = 8.6%**

| Feature Name | Feature Type | Loading |
|---|---|---|
| NucCVDisc3 | Nucleus texture | 0.1919 |
| NucCVDisc4 | Nucleus texture | 0.1902 |
| NucCVCentroidDist | Nucleus morphometry | 0.1825 |
| NucFracTotalDisc1 | Nucleus texture | 0.1786 |
| NucVarDisc3 | Nucleus texture | 0.1745 |
| NucRatioMajorMinor | Nucleus morphometry | 0.1741 |
| NucCircularity | Nucleus morphometry | -0.1726 |
| NucCVDisc2 | Nucleus texture | 0.1693 |
| NucFracTotalDisc2 | Nucleus texture | 0.1658 |
| NucVarDisc2 | Nucleus texture | 0.1652 |

**Movie S1. Fluorescence time lapse of an untreated WT MEF over a 60h time course.**

**Movie S2. Fluorescence time lapse of an Apoptosis[+] cell over a 64h time course.**

**Movie S3. Fluorescence time lapse of an Apoptosis[-] cell over a 64h time course.**
